# Mammalian returns to the sea reveal broad genomic slowing rather than a fixed adaptive toolkit

**DOI:** 10.64898/2026.08.26.747444

**Authors:** Jiaqi Wu, Takahiro Yonezawa, Naoki Kohno, Hirohisa Kishino

## Abstract

Marine mammals — cetaceans, pinnipeds, sirenians, sea otters and polar bears — returned to the sea independently, yet whether their genomes converged on a shared adaptive programme or shifted in a common direction without a fixed toolkit has remained unclear. Here we separate marine specialization from general aquatic dependence across 302 mammals and 17,432 protein-coding genes and show that the dominant genomic signature of marine life is widespread evolutionary slowing, not acceleration: of 1,559 marine-associated genes, nearly 88% evolved more slowly, and this slow-direction bias persisted (98%) after removing cetaceans. Compact gene fingerprints that distinguish marine identity combine fast-rate remodeling of body-surface and sensory genes with slow-rate constraint on blood, metabolic and DNA-repair genes, but these fingerprints are sharpened by cetaceans and do not preserve a fixed functional toolkit across lineages. Species-level and ancestral-branch decompositions reveal that different marine mammals assembled marine-like genomic states through distinct gene combinations. Mammalian marine convergence is therefore directional rather than modular: a broad constraint landscape resolved into clade-weighted genomic fingerprints.

## 1. Introduction

The repeated return of mammals to the sea is a textbook example of convergent evolution. Cetaceans, pinnipeds, sirenians, sea otters and polar bears all entered marine environments from terrestrial ancestors, facing overlapping demands on diving, thermoregulation, oxygen transport, salt and water balance, body-surface biology and sensory systems ^1–3^. Much of comparative genomics has therefore searched for a shared adaptive signature behind these transitions: recurrent amino-acid substitutions, accelerated evolution, or repeated targeting of the same biological pathways.

Previous studies have found important pieces of this signal. Genome-wide analyses reported parallel amino-acid substitutions and convergent changes in genes related to skin, hearing and other marine-associated traits ^4,5^. Broader sampling of aquatic mammals identified candidate loci associated with diving physiology, immunity and thermoregulation ^6^. Rate-based analyses added a different view, showing that convergence can also appear as coordinated shifts in evolutionary constraint across many genes, including widespread deceleration rather than only recurrent amino-acid change ^7^. SplitAligner enabled gene-wise branch lengths to be projected onto a common species-tree coordinate system ^8^, and the gene–branch interaction framework provided a way to quantify gene-specific rate deviations across branches ^9^. Together, these studies established that marine life leaves detectable genomic traces, but they did not resolve what kind of convergence dominates the mammalian return to the sea.

Three distinctions are critical. First, marine specialization is not the same as general aquatic dependence: hippopotamuses and river otters are strongly aquatic but non-marine, whereas polar bears and sea otters are marine-associated but far less aquatic than dolphins. Pooling these taxa into a single “aquatic” or “marine” signal risks confounding distinct ecological pressures. Second, adaptive innovation at individual candidate genes does not establish whether the dominant genome-wide signal is acceleration or slowing — a question that has not been tested at scale while cleanly separating marine from non-marine aquatic lineages. Third, the genes that best distinguish marine mammals from other species need not be the same as the genes that shift most consistently across lineages. Molecular convergence may therefore take the form of a fixed adaptive toolkit — the same genes and pathways deployed repeatedly — or it may instead be directional: many genes shifting in the same broad direction while different lineages assemble distinct genomic fingerprints.

Here we test these alternatives across 302 mammals and 17,432 protein-coding genes by separating marine specialization from independently scored general aquatic dependence. We show that the dominant genomic signature of marine life is widespread evolutionary slowing, not acceleration: 1,366 of 1,559 marine-associated genes evolved more slowly, and after removing cetaceans, 894 genes remained significant with 98% in the same slow direction. This broad slowing is shared with general aquatic dependence and is enriched for hematological and systemic physiology. In the compact fingerprints, the return-to-water signal is resolved into physiological systems that combine fast-rate remodeling of external interfaces with slow-rate constraint on internal functions such as blood regulation, metabolism, DNA repair and reproductive-cell biology. Yet the compact genomic fingerprint that distinguishes marine identity is more lineage-weighted: it is sharpened by cetaceans, reuses some individual genes across comparisons, but does not preserve a fixed functional toolkit. Species-level and ancestral-branch profiles further reveal that different marine mammals assemble marine-like genomic states through distinct gene combinations. Mammalian returns to the sea therefore reveal directional, not modular, genomic convergence: a broad constraint landscape resolved into clade-weighted biological fingerprints.

## 2. Results

### 2.1 Marine specialization is distinct from general aquatic dependence

Not all aquatic mammals are marine, and not all marine mammals are strongly aquatic. The hippopotamus spends much of its life in water yet inhabits freshwater rivers and lakes; it is aquatic but not marine. The polar bear depends on the marine environment for hunting and survival, yet it walks on sea ice and land — ecologically marine, but far less aquatic than a dolphin. River dolphins belong to cetacean lineages that evolved in the ocean, yet they now live exclusively in freshwater. These cases show that marine specialization and general aquatic dependence are not interchangeable, and any genomic analysis that conflates them risks missing the distinct pressures each imposes.

We therefore adopted a two-layer trait framework (Fig. 1). Marine membership was assigned using established marine mammal authorities, yielding 51 marine species. General aquatic dependence was quantified independently using a five-dimensional scoring system encompassing foraging medium, locomotion, reproduction, morpho-physiological specialization, and aquatic time budget, with summed scores ranging from 0 to 18 (see Methods). This scoring classified the remaining non-marine species into 12 non-marine aquatic mammals (such as hippopotamus, beaver, and river otters), 13 semi-aquatic mammals, and 226 terrestrial species.

**Fig. 1.**
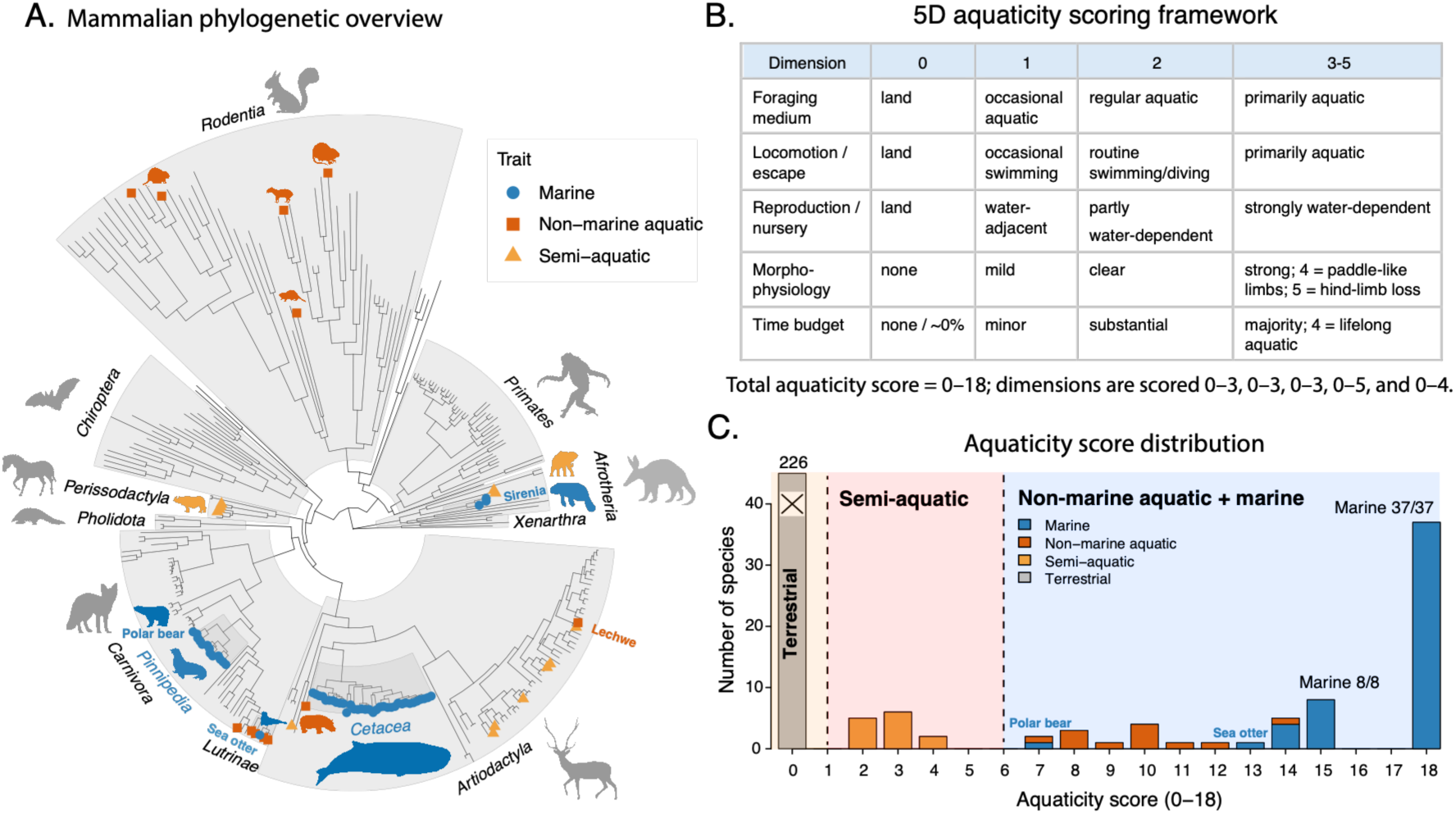
Marine specialization and general aquatic dependence are distinct ecological dimensions across mammals. (A) Phylogenetic distribution of marine (blue circles), non-marine aquatic (orange squares) and semi-aquatic (yellow triangles) mammals across the 302-species dataset. Marine membership follows established marine mammal authorities; non-marine aquatic and semi-aquatic categories were defined using an independently curated aquaticity framework. (B) Five-dimensional aquaticity scoring framework used to quantify general aquatic dependence, spanning foraging medium, locomotion or escape, reproduction or nursery, morpho-physiological specialization and aquatic time budget. The total aquaticity score ranges from 0 to 18; full scoring criteria are provided in Supplementary Table S1. (C) Distribution of aquaticity scores across the 302 species. Of these, 226 scored zero; the remaining 76 spanned semi-aquatic, non-marine aquatic and marine states. No sampled species scored 5 or 6, yielding an empirical gap used to operationalize the semi-aquatic/aquatic boundary. Marine mammals occupied the highest scores, but high aquaticity alone did not define marine identity: the polar bear (score 7) and sea otter (score 13) were classified as marine despite scoring below fully aquatic cetaceans.

Principal component analysis confirmed that the five scoring dimensions collapsed onto a single dominant gradient of aquatic dependence (PC1 explained 97.0% of variance, Fig. S1), validating the summed aquaticity score as a compact quantitative axis. But aquaticity score alone did not determine marine identity: the polar bear scored 7, the sea otter scored 13, and fully marine cetaceans scored 18, yet all three were classified as marine. Marine specialization is therefore nested within general aquatic dependence but is not simply its extreme end.

For downstream analyses, marine status was modelled as a binary trait (marine versus all non-marine species). General aquatic dependence was modelled as a separate binary trait contrasting strongly aquatic species (marine plus non-marine aquatic, n = 63) against terrestrial species (n = 226), with the 13 semi-aquatic species excluded to preserve an unambiguous two-state contrast.

### 2.2 Evolutionary rate profiles predict marine and aquatic identity

Can the evolutionary history encoded in protein-coding genes tell us whether a mammal lives in the ocean? We addressed this by quantifying how each gene’s evolutionary rate deviates from its genome-wide background on each branch of the mammalian tree — a measure called the gene–branch interaction (GBI; see Methods). GBI captures gene-specific rate shifts that cannot be explained by overall mutation rate or divergence time differences, and thereby isolates lineage-specific changes in functional constraint or selective pressure.

We computed GBI values for 17,432 protein-coding genes across 302 mammals and used penalized logistic regression (LASSO) to ask whether small subsets of genes could distinguish marine or aquatic mammals from other species. To guard against overfitting, we evaluated predictive accuracy by genus-level leave-one-out cross-validation, repeating gene selection within each training fold so that held-out species never influenced which genes entered the model (see Methods).

Under this rigorous validation, evolutionary rate profiles robustly predicted marine identity (AUC = 0.936). Prediction of general aquatic dependence was weaker but still significant (AUC = 0.826; Fig. 2B). Marine specialization thus left a sharper signature in gene-level evolutionary rates than did general aquatic dependence alone, consistent with the stronger and more uniform ecological demands — diving, thermoregulation, osmotic balance and sensory remodeling — that characterize marine life.

**Fig. 2.**
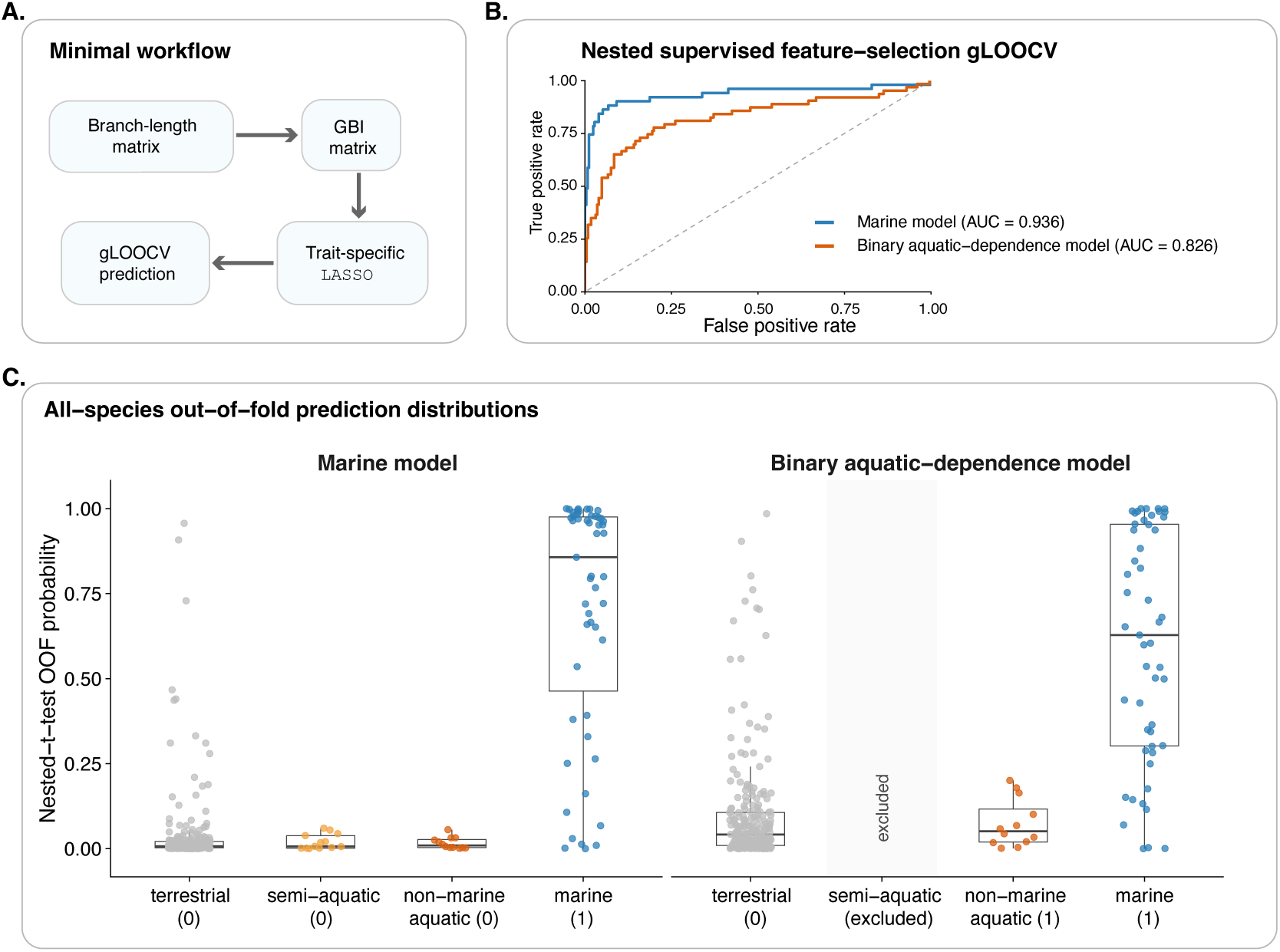
Evolutionary rate profiles distinguish marine identity more sharply than general aquatic dependence. (A) Workflow overview. For each gene, branch-specific evolutionary rates were expressed as gene–branch interaction (GBI) values that capture deviations from the genome-wide background rate on each branch. GBI profiles were then used to ask whether gene-rate patterns distinguish marine specialization or aquatic dependence; the sparse gene-fingerprint model was implemented with penalized logistic regression (LASSO). (B) Held-out discrimination under genus-level leave-one-out cross-validation, with gene selection repeated inside each training fold so that held-out species did not influence gene choice. The marine model achieved an AUC of 0.936; the aquatic-dependence model achieved 0.826. (C) Distribution of held-out profile scores across ecological categories from the same cross-validation procedure. The marine model separated marine taxa from all other categories at the aggregate level. The aquatic-dependence model showed a broader distribution, with the strongest signal concentrated among marine-positive taxa. Semi-aquatic species were excluded from aquatic-dependence training and AUC calculation and are annotated as excluded.

To visualize this predictive signal, we plotted the distribution of out-of-fold probabilities across species grouped by ecological category (Fig. 2C). The marine model separated marine taxa from terrestrial, semi-aquatic and non-marine aquatic species at the aggregate level. The aquatic-dependence model showed a broader distribution, with the strongest signals concentrated among marine mammals rather than being uniformly recovered across all aquatic taxa.

### 2.3 Return to the sea is marked by widespread evolutionary slowing

Having established that gene-level rate profiles can predict marine identity, we next asked what direction these rate changes take. Do genes evolve faster in marine lineages, reflecting a burst of adaptive innovation, or slower, reflecting tightened functional constraint?

The answer was overwhelmingly in one direction. Across 17,258 genes tested, 1,559 showed significantly different evolutionary rates on marine branches (FDR ≤ 0.01). Of these, 1,366 — nearly 88% — evolved more slowly in marine mammals, whereas only 193 evolved faster (Fig. 3A). This striking asymmetry indicates that the dominant genomic signature of marine adaptation is not accelerated evolution but widespread strengthening of functional constraint. Genes that slow down in marine lineages are enriched for functions in blood physiology, including erythrocyte and hemoglobin regulation, platelet biology, and cardiovascular function, as well as body size and systemic metabolic traits (Fig. 3C).

**Fig. 3.**
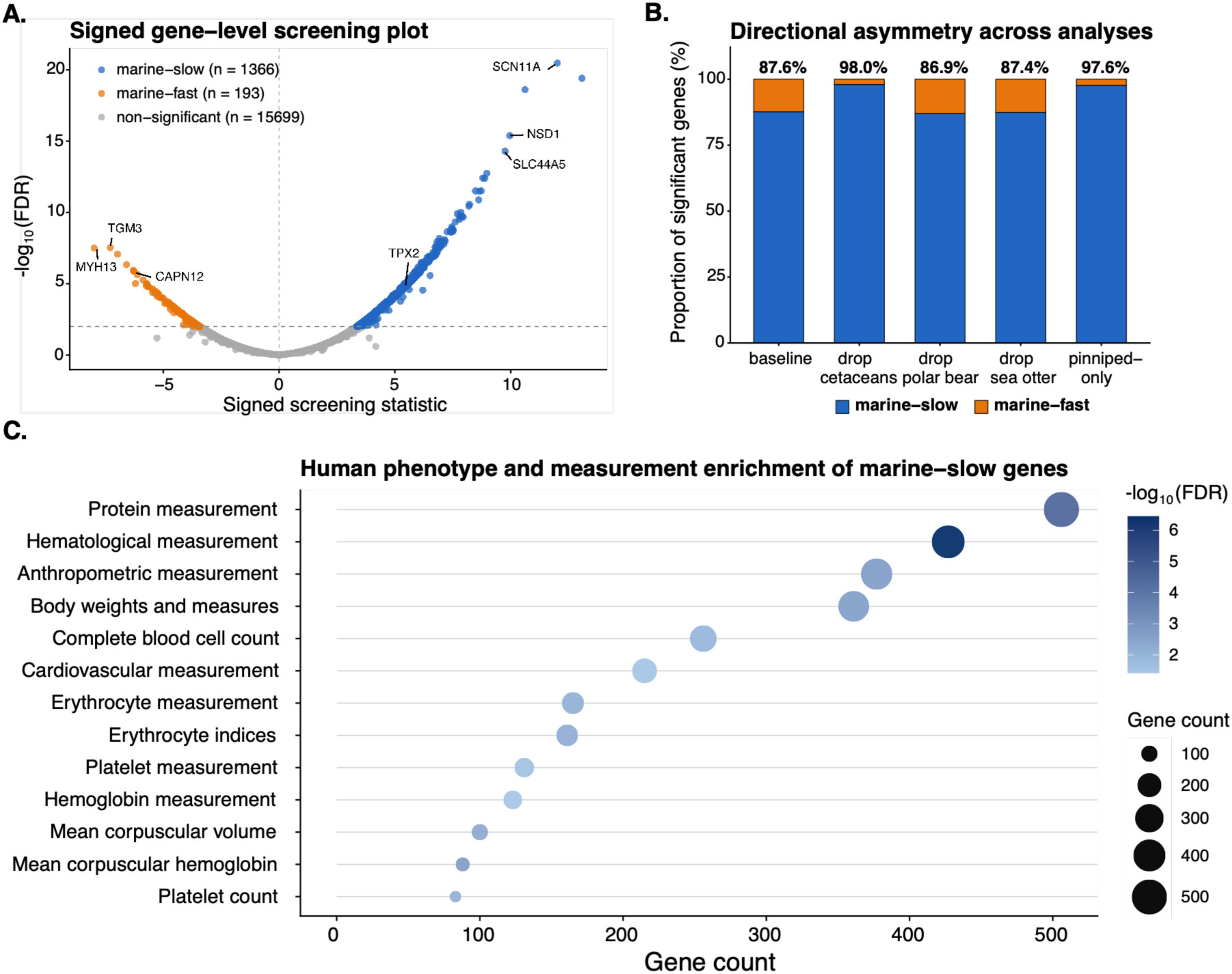
Return to the sea is dominated by evolutionary slowing, not acceleration. (A) Genome-wide screening of individual genes for rate shifts associated with marine life. Of 17,258 genes tested, 1,559 were significant at FDR ≤ 0.01. Of these, 1,366 evolved more slowly on marine branches (marine-slow; blue) and only 193 evolved faster (marine-fast; orange), indicating that nearly 88% of marine-associated genes evolved in the direction expected under strengthened constraint. The x-axis shows the signed screening statistic: positive values indicate slower evolution in marine lineages. (B) This slow-direction dominance was robust across sensitivity analyses. After removing cetaceans, 98.0% of significant genes remained slow-direction; similar proportions held after removing the polar bear, the sea otter, or restricting to pinnipeds alone. (C) Functional enrichment of the 1,366 marine-slow genes. Point size indicates gene count; colour indicates enrichment significance (−log₁₀ FDR). Marine-slow genes were enriched for blood physiology, including erythrocyte, hemoglobin, platelet, cardiovascular and body-measurement terms.

This slow-direction pattern was not a peculiarity of a particular clade. When we repeated the screen after removing cetaceans, 894 genes remained significant, and 98.0% of them were in the slow direction (Fig. 3B). Similar results held after removing the polar bear, the sea otter, or restricting the analysis to pinnipeds alone (86.9–98.0% slow across all sensitivity runs). The constraint signal is therefore broadly shared across independently evolved marine lineages.

General aquatic dependence showed a parallel pattern: 1,055 of 1,227 significant genes (86.0%) evolved more slowly on aquatic-associated branches. The marine and aquatic slow-gene sets overlapped extensively — 983 genes were significant and slow-direction in both screens, accounting for 72.0% of marine-slow genes and 93.2% of aquatic-slow genes. Among genes significant in both comparisons, directional assignments were fully concordant: every gene that slowed in marine lineages also slowed in aquatic lineages, and vice versa.

This result reconciles with previous reports of accelerated evolution in specific marine adaptation genes, because the two patterns operate at different levels. Individual candidate genes may undergo rapid amino acid substitution driven by positive selection, while the broader genomic background simultaneously tightens constraint. Diving, thermoregulation, osmotic balance, and oxygen transport all impose functional requirements that limit how far molecular systems can drift — producing the widespread slowdown detected here. Targeted acceleration in specific genes and broad constraint across many genes are not mutually exclusive; they coexist as complementary layers of marine genome evolution.

### 2.4 Compact genomic fingerprints combine remodeling and constraint across physiological systems

After establishing predictive performance through cross-validation, we fitted final models to the complete dataset to summarize which genes contributed to marine and aquatic-dependence prediction and in which direction. The marine model retained 71 predictors and the aquatic-dependence model retained 101; 24 predictors were shared, whereas 47 were marine-specific and 77 were aquatic-specific (Fig. 4A). These predictor sets overlapped but were far from interchangeable — consistent with the ecological distinction between marine specialization and general aquatic dependence.

**Fig. 4.**
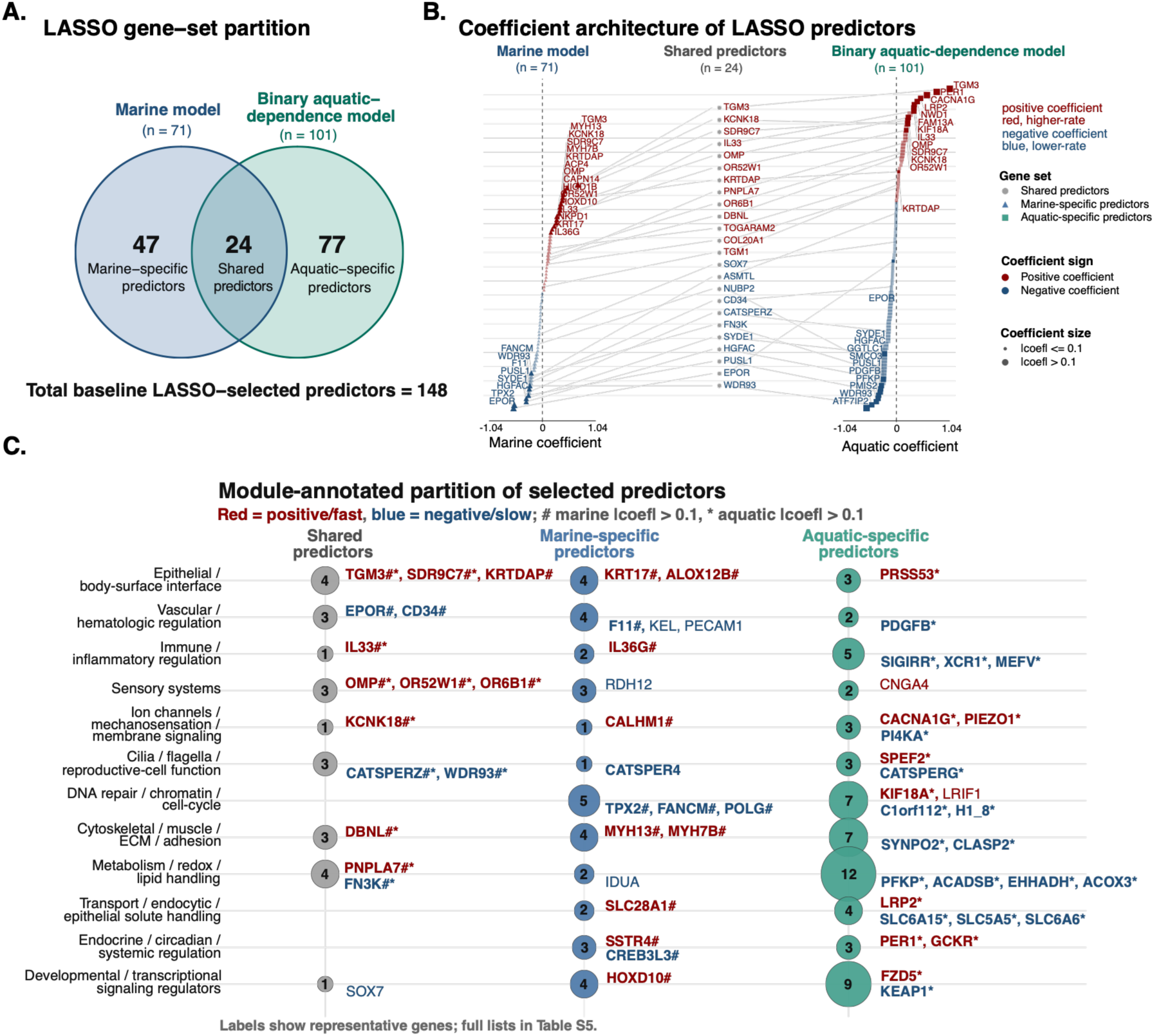
Compact marine and aquatic fingerprints combine fast-rate and slow-rate components across physiological systems. After validation, final full-data gene fingerprints summarize the retained genes and directional contributions underlying marine specialization and aquatic dependence; cross-validated performance is reported separately in Figs. 2 and 5A. (A) Gene-set partition. The marine fingerprint retained 71 genes and the aquatic-dependence fingerprint 101, with 24 shared, 47 marine-specific and 77 aquatic-specific, totalling 148 unique genes. (B) Coefficient structure. For each gene, the coefficient sign indicates whether higher evolutionary rates (red; fast-side, consistent with remodeling or relaxed constraint) or lower rates (blue; slow-side, consistent with strengthened constraint) contribute to the marine or aquatic score. Coefficient sign describes contribution direction within the fitted fingerprint; single-gene slow/fast evidence is reported separately in Fig. 3 and Fig. S2. Shared genes carry both fingerprint coefficients; trait-specific genes appear only on one side. Larger points mark |coefficient| > 0.1. (C) Functional organization across 12 curated display categories. Circle size indicates the number of genes per cell. Representative genes are labelled for readability; hash marks indicate |marine coefficient| > 0.1, asterisks indicate |aquatic-dependence coefficient| > 0.1. Body-surface, sensory and ion-channel genes are predominantly fast-side, whereas blood, DNA-repair and metabolic genes are predominantly slow-side. This grouping is descriptive and does not constitute formal evidence for recurrent functional modules, which is tested in Fig. 5C. Full gene lists are in Supplementary Table S5.

Importantly, the selected predictors were not uniformly slow or fast. In the LASSO models, each predictor carries a coefficient whose sign indicates whether higher or lower relative evolutionary rates contribute to the trait score (see Methods). We refer to predictors with positive coefficients as fast-side components (higher relative rates push the score toward the marine or aquatic state) and those with negative coefficients as slow-side components (lower relative rates push the score toward the trait state). The sparse fingerprints therefore combine both directions of rate change, and the biological systems they represent fall into a striking pattern (Fig. 4B, C). These directional labels describe model-score contributions within the fitted sparse architecture; genome-wide single-gene rate direction is analysed separately in Fig. 3 and Fig. S2.

#### Genes facing the external environment are predominantly fast-side

Body-surface and epithelial predictors were almost entirely fast-side components: the transglutaminases TGM3, TGM1 and TGM5 ^10,11^, the epidermal lipid enzyme SDR9C7 ^12^, the keratinocyte marker KRTDAP ^13^, the lipid-processing enzyme ALOX12B ^14^, and the keratins KRT17 ^15^ and CAPN14 ^16^ all carried positive coefficients. This pattern is consistent with remodeling of the skin barrier during the transition to aquatic life — a shift from fur-dominated insulation toward blubber-assisted thermoregulation and modified epidermal lipid-barrier architecture. Sensory-system predictors showed a parallel pattern: the olfactory genes OMP, OR52W1 and OR6B1 ^17,18^, the cone photoreceptor phosphodiesterase PDE6C ^19^, and the cyclic-nucleotide-gated channel CNGA4 ^20^ were all fast-side, consistent with the well-documented regression of olfactory and visual pathways in aquatic mammals. Ion-channel and mechanosensory predictors reinforced this trend: KCNK18 ^21^ (pain perception), CACNA1G ^22^ (T-type calcium channel), PIEZO1 ^23^ (mechanosensation) and CALHM1 ^24^ (calcium homeostasis) were all fast-side, pointing to broad remodeling of sensory and membrane-signaling thresholds.

#### Genes governing internal physiology are predominantly slow-side

Vascular and hematologic predictors showed the opposite pattern: the erythropoietin receptor EPOR ^25^, the hepatocyte growth factor activator HGFAC ^26^, the hematopoietic marker CD34 ^27^, and the coagulation factor F11 all carried negative coefficients, indicating that lower relative evolutionary rates at these genes contributed to marine and aquatic scores. Additional slow-side vascular predictors included the platelet glycoprotein ITGA2B ^28^ and the growth factor PDGFB ^29^— all consistent with strengthened constraint on blood-oxygen transport, hemostasis and vascular integrity in diving mammals. A single vascular gene, HIGD1B ^30^ (a hypoxia-responsive mitochondrial regulator), was marine-specific and fast-side, suggesting a distinct role in oxygen-sensing rather than the broader constraint pattern. DNA-repair and chromatin-maintenance predictors were likewise predominantly slow-side: TPX2 ^31^ (spindle assembly and DNA-damage response), ATF7IP2 ^32^ (chromatin regulation), FANCM ^33^ (Fanconi anaemia DNA-repair pathway), POLG ^34^ (mitochondrial DNA polymerase) and the linker histone H1.8 all carried negative coefficients. These slow-side signals point to strengthened maintenance of genome integrity in species that face elevated oxidative stress from repeated diving and prolonged apnea. Metabolic predictors followed the same direction: the glycolytic regulator PFKP ^35^, the fatty-acid oxidation enzymes ACADSB ^36^ and EHHADH ^37^, the peroxisomal oxidase ACOX3 ^38^, and the glycation-protective enzyme FN3K ^39^ were all slow-side, suggesting tightened constraint on energy metabolism and redox homeostasis. Reproductive-cell and ciliary predictors were directionally split. The sperm calcium-channel components CATSPERZ, CATSPERG and CATSPER4 ^40^, together with the ciliary factor WDR93 ^41^, lay on the slow-side, consistent with conserved gamete and ciliary function under aquatic conditions. However, the ciliary-motility gene SPEF2 ^42^, the centriole-biogenesis factor DEUP1 ^43^, and the ciliary microtubule protein TOGARAM2 ^44^ were fast-side, suggesting that some components of ciliary biology are being remodeled rather than merely conserved.

#### Circadian and immune predictors show mixed directionality

Not all functional categories fell neatly into one direction. The circadian clock gene PER1 ^45^ carried the largest single positive coefficient in the aquatic-dependence model (+0.784), placing it strongly on the fast-side, while the melatonin-pathway-adjacent ASMTL ^46^ and the metabolic regulator CREB3L3 ^47^ were slow-side. Among immune predictors, the interleukins IL33 and IL36G ^48^ were fast-side, while the inflammasome component NLRP8 ^49^, the immune regulators SIGIRR and XCR1 ^50^, and the innate-immunity gene MEFV ^51^ were slow-side. This directional heterogeneity suggests that immune and circadian systems are not uniformly remodeled or constrained during aquatic transitions but are instead reshaped in gene-specific ways.

Taken together, the sparse predictor architectures reveal a directional logic: the transition to aquatic and marine life combines relaxation-like rate shifts at genes mediating external interfaces — skin, sensation, ion-channel signaling — with constraint-like rate shifts at genes governing internal physiology — blood, DNA repair, metabolism and reproductive-cell function. This directional split provides a more informative summary of the genomic fingerprints than gene identity alone, and it connects the broad slow-rate signal identified in Section 2.3 to specific physiological systems. We note that this functional organization is descriptive; whether these predictor categories recur stably across lineage-removal analyses is evaluated separately below.

### 2.5 Cetaceans sharpen the compact marine fingerprint, but broad slowing spans marine lineages

Whales and dolphins are the most species-rich and deeply aquatic marine mammals in our dataset, raising a central question: are we detecting a general marine-mammal signal, or mostly a cetacean signal? To separate these possibilities, we removed or isolated major marine lineages and compared two layers of evidence: the genome-wide slow-rate signal and the compact genomic fingerprints used for prediction (Fig. 5).

**Fig. 5.**
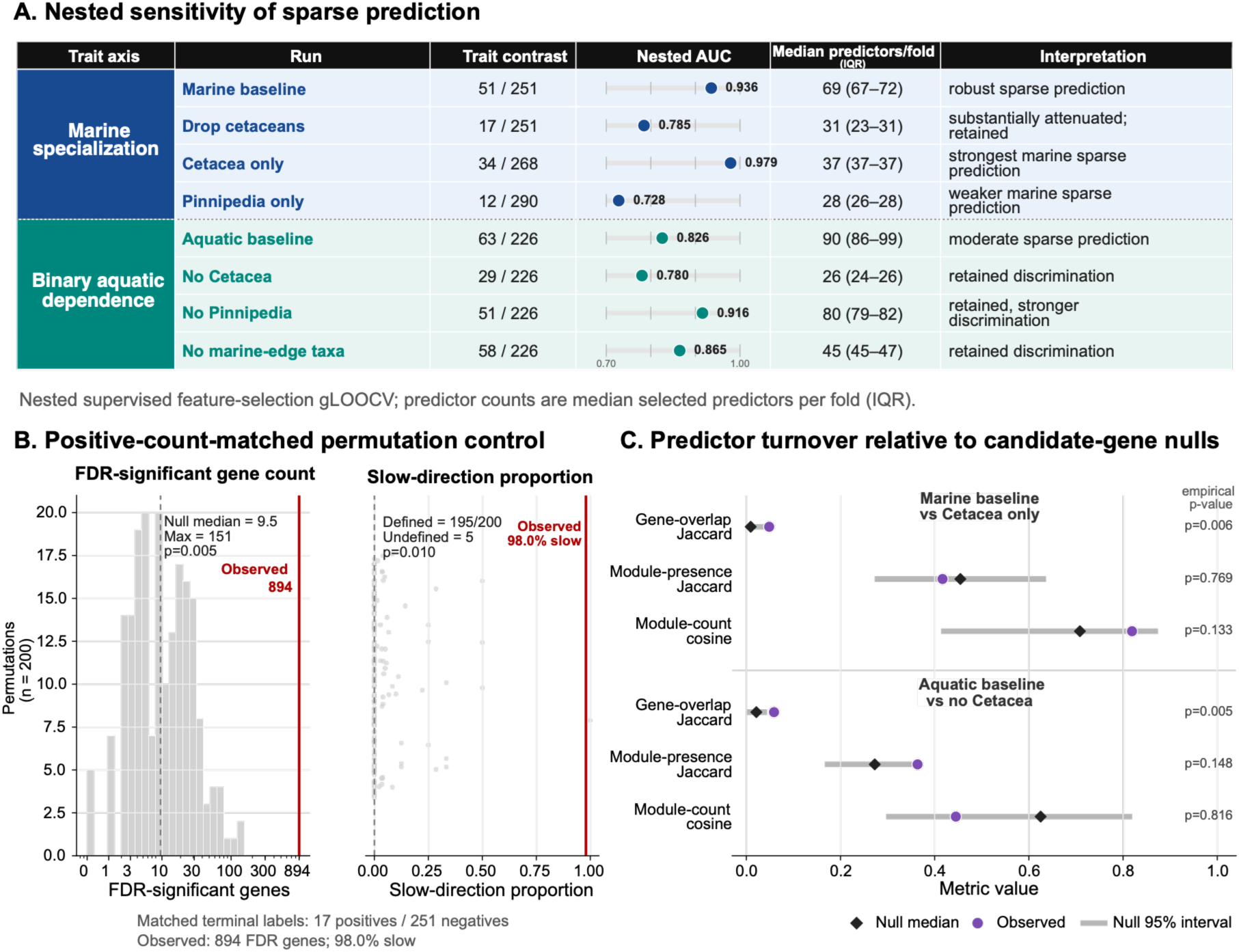
Cetaceans sharpen the compact marine fingerprint, but broad slowing persists without them. (A) Lineage-removal tests of the compact marine and aquatic-dependence fingerprints, using the same held-out cross-validation as Fig. 2. Marine discrimination was strongest in the full dataset and in the cetacean-only contrast (AUC 0.936 and 0.979) and was substantially weakened by removing cetaceans (0.785) or restricting to pinnipeds (0.728). Aquatic-dependence discrimination was weaker overall (0.826) but was less dependent on any single clade. (B) Broad slowing persists without cetaceans. After removing all cetaceans, 894 genes remained significant, of which 876 (98.0%) were slow-direction — exceeding expectations from positive-count-matched permutations (P_count = 0.005; P_slow = 0.010). (C) Recurring genes, not fixed modules. Gene-level overlap between fingerprints from different lineage contrasts exceeded null expectations from the available trait-associated gene pool, but similarity at the level of curated functional modules did not. The compact fingerprints therefore reuse some genes more than expected by chance, without reducing to a fixed functional toolkit. Numerical values supporting A–C are in Supplementary Table S6.

The broad slow-rate signal was not a cetacean-only pattern. After removing all cetaceans, the marine single-gene screen still recovered 894 significant genes at FDR ≤ 0.01, of which 876 were slow-direction genes — meaning 98.0% of the significant genes outside cetaceans still evolved more slowly on marine-associated branches. Positive-count-matched permutations confirmed that this result was not explained simply by the smaller number of remaining marine-positive species (P_count = 0.005; P_slow = 0.010; Fig. 5B). Pinnipeds, sirenians, the polar bear and the sea otter therefore retain a strong constraint-associated rate signal independently of whales and dolphins.

The compact marine fingerprint behaved differently. Marine identity was most sharply recoverable in the full baseline dataset and in the cetacean-only contrast, with cross-validated AUC values of 0.936 and 0.979, respectively (Fig. 5A). Removing cetaceans substantially weakened, but did not eliminate, the signal (AUC = 0.785), and restricting the positive class to pinnipeds alone yielded weaker discrimination (AUC = 0.728). Thus, cetaceans do not create the broad slow-rate background, but they strongly sharpen the small-gene fingerprint that distinguishes marine identity — likely reflecting their long evolutionary history in water, deep physiological commitment to aquatic life, and comparatively rich taxonomic sampling in our dataset.

General aquatic dependence showed a different lineage structure. Its baseline fingerprint was weaker than the marine model (AUC = 0.826), but it was less dominated by any single marine clade: discrimination was retained after removing cetaceans (AUC = 0.780), removing pinnipeds (AUC = 0.916), or excluding marine-edge taxa such as sirenians, the polar bear and the sea otter (AUC = 0.865). This pattern reinforces the two-layer trait framework: aquatic dependence is broader and more heterogeneous than marine specialization, whereas marine specialization is more sharply compressed by cetacean-rich rate profiles.

We next asked whether the compact fingerprints preserved the same genes or the same functional categories across lineage-removal analyses. Gene-level overlap between corresponding predictor sets exceeded comparison-specific null expectations for both the marine baseline-versus-cetacean-only and aquatic baseline-versus-no-cetacean comparisons (Fig. 5C). However, similarity at the level of curated functional modules did not exceed the corresponding null expectations. The compact fingerprints therefore reuse some genes more than expected by chance, but they do not reduce to a fixed catalogue of recurrent functional modules — consistent with selection acting on individual genes within a wide functional landscape rather than on conserved pathways.

Together, these results reveal that marine genomic convergence is directional rather than modular. A broad, slow-direction genomic background spans multiple marine lineages and persists without cetaceans — the hallmark of shared constraint, not a shared adaptive toolkit. Over this background, cetaceans sharpen a narrower compact fingerprint that allows marine identity to be distinguished from small gene sets, but this fingerprint is clade-weighted rather than universal. The convergent signal is therefore real, but it lives in the direction of evolutionary change, not in a fixed catalogue of genes or pathways.

### 2.6 Species and ancestral fingerprints reveal distinct genomic routes back to the sea

The analyses above describe the marine signal in aggregate: broad evolutionary slowing, compact gene fingerprints, and cetacean sharpening. But marine mammals are not an aggregate — each lineage returned to water from a different starting point and assembled a different genomic profile. We therefore used the final marine and aquatic fingerprints as reference maps for representative living species and selected ancestral branches, and decomposed each profile into gene-by-gene contributions (Fig. 6). These profiles show how individual lineages combine the shared gene vocabulary into distinct marine-like and aquatic-like genomic fingerprints.

**Fig. 6.**
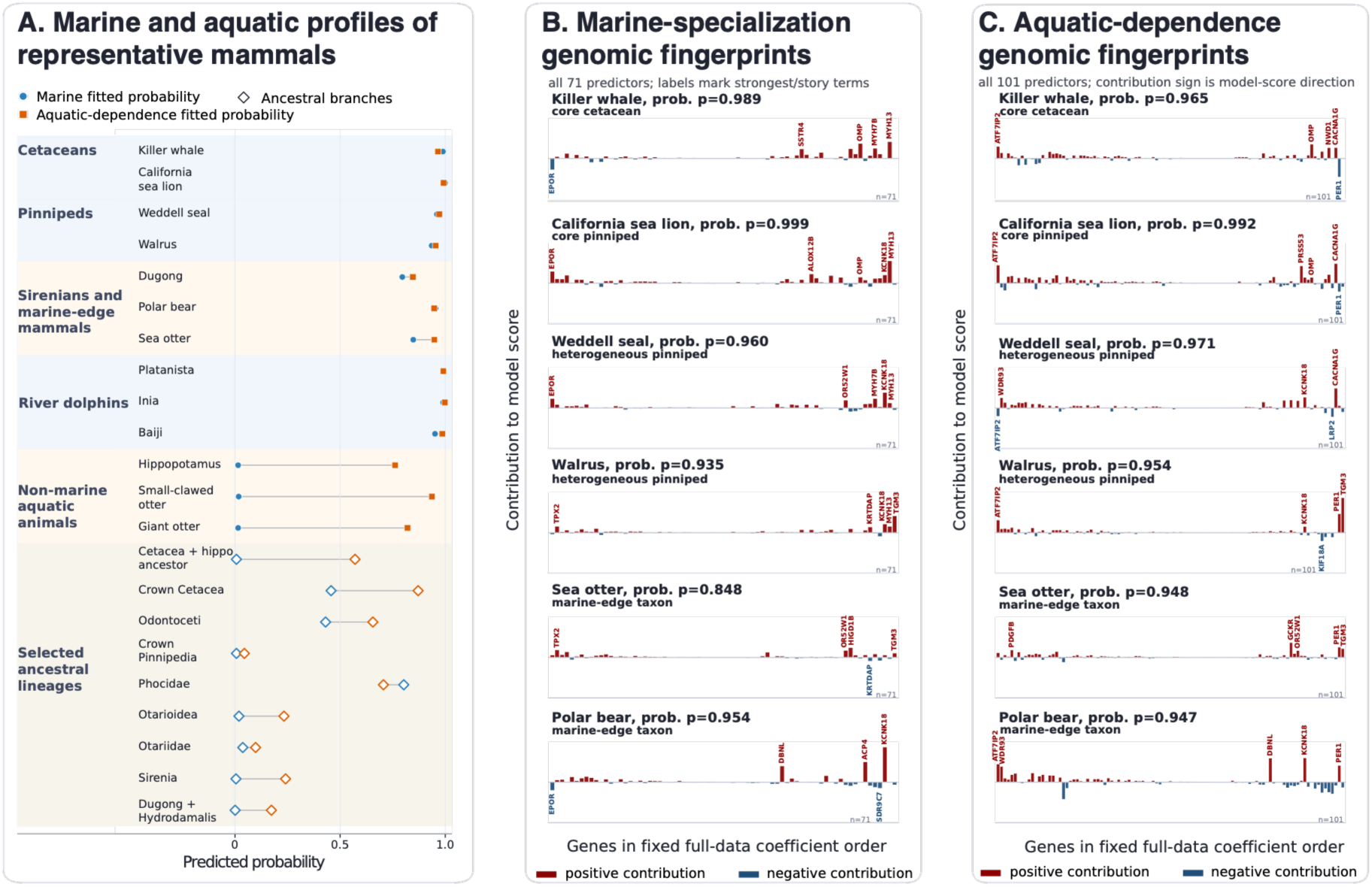
Species and ancestral fingerprints reveal distinct genomic routes back to the sea. (A) Representative terminal species and selected ancestral branches mapped onto the marine-specialization and aquatic-dependence axes. Filled symbols show terminal species; open diamonds show ancestral branches. Marine-like and aquatic-dependence profile scores were computed independently by projecting each species or branch onto the final gene fingerprints defined after validation; a species can therefore score high on one axis and low on the other. Ancestral-branch scores are rate-based genomic profiles and should not be read as direct habitat assignments. (B) Marine-specialization genomic fingerprints for six representative species. Each bar represents the contribution of one of the 71 marine-fingerprint genes, plotted in fixed coefficient order. Contributions were calculated as scaled GBI × coefficient; the intercept is omitted. Red bars push the profile score toward the marine state, whereas blue bars push it away. Gene labels mark the strongest contributors and genes discussed in the text. Annotations below each species name indicate ecological grouping: core cetacean, core pinniped, heterogeneous pinniped or marine-edge taxon. (C) Aquatic-dependence genomic fingerprints for the same six species, using the 101 aquatic-dependence genes. Layout and colour coding follow panel B. Held-out discrimination is reported in Figs. 2 and 5A; the fingerprints shown here are descriptive projections onto the final gene fingerprints after validation.

#### Several genes appear as prominent contributors across multiple lineages

Although Section 2.5 established that no fixed functional module recurs across lineage-removal analyses, the species-level decompositions revealed that certain individual genes contributed prominently and repeatedly to marine-like profiles (Fig. 6B). MYH13 ^52^, a superfast myosin expressed in extraocular and laryngeal muscles, was among the strongest positive contributors in the killer whale, California sea lion, Weddell seal and walrus — spanning both cetaceans and pinnipeds. KCNK18 ^21^, a potassium channel linked to pain perception, contributed to the marine scores of the California sea lion, Weddell seal, walrus and polar bear. EPOR, the erythropoietin receptor ^25^, contributed across the killer whale, California sea lion, Weddell seal and polar bear in the slow-rate direction — consistent with tightened constraint on oxygen-carrying capacity. These recurrent individual genes do not constitute a shared adaptive toolkit (their module-level context does not recur above null expectations), but they suggest that independently evolved marine lineages have faced overlapping functional demands on muscle physiology, sensory modulation and blood-oxygen transport, even though the broader pathway context differs among lineages.

#### Each species assembles a distinct marine-like fingerprint

The killer whale (marine-like profile 0.989) drew prominently on SSTR4 ^53^, a somatostatin receptor involved in nociception, and OMP ^18^, an olfactory marker protein — neither of which appeared among top contributors in pinnipeds. The California sea lion (0.999) instead drew on ALOX12B ^54^, an epidermal lipid-processing enzyme absent from cetacean fingerprints. These differences reflect the distinct body plans and ecological strategies of fully aquatic cetaceans and amphibious pinnipeds: cetacean fingerprints emphasize sensory regression and deep-diving physiology, whereas pinniped fingerprints draw more on epithelial and integumentary remodeling.

#### Pinnipeds show heterogeneous fingerprints

The California sea lion formed the cleanest pinniped marine profile, annotated in Fig. 6 as a “core pinniped.” The Weddell seal and walrus, annotated as “heterogeneous pinnipeds,” assembled their marine-like scores from different gene combinations. The walrus (0.935) drew more heavily on skin-barrier predictors (KRTDAP, TGM3) and TPX2 ^31^, a cell-division regulator, whereas the Weddell seal (0.960) featured OR52W1, MYH7B and KCNK18 more prominently. This within-group heterogeneity is consistent with the ancestral-branch analysis: crown Pinnipedia showed low marine-like and aquatic-like genomic profile scores, suggesting that the strong marine fingerprints of extant pinnipeds were assembled after — not before — the divergence of major pinniped subclades.

#### Marine-edge taxa decouple ecological classification from genomic profiles

The polar bear and sea otter are classified as marine mammals based on their ecological dependence on the ocean, yet their genomic fingerprints look different from those of cetaceans and pinnipeds. The polar bear (marine-like profile 0.954) drew on a largely distinct gene set — DBNL ^55^ (cytoskeletal organization), ACP4 ^56^(mineralized tissue), SDR9C7 ^12^ (lipid metabolism) — with minimal overlap with cetacean contributors. Among the six extant species decomposed in Fig. 6B, the sea otter showed the weakest marine-like profile (0.848), with its strongest contributors coming from skin-barrier genes (KRTDAP, TGM3) and HIGD1B (hypoxia response). In the aquatic-dependence fingerprint, the sea otter scored notably higher than on the marine axis (0.948 vs 0.848), whereas the polar bear scored similarly high on both axes but with a distinctly different gene composition from cetaceans and pinnipeds. Both taxa thus illustrate marine-edge genomic profiles: they are ecologically marine but do not share the deeply aquatic genomic fingerprints of whales and seals.

#### The circadian clock gene PER1 shows lineage-dependent directionality

In the aquatic-dependence fingerprints (Fig. 6C), PER1 ^45^ appeared among the top contributors across most species — but with a notable exception. In the killer whale, PER1 contributed on the constrained side of the aquatic fingerprint, whereas in pinnipeds, the polar bear and the sea otter it contributed on the remodeling side. This divergence may reflect different light environments experienced by fully pelagic deep-diving cetaceans and ice-associated or coastal species, but this functional interpretation remains hypothesis-generating.

#### River dolphins retain marine-like profiles to varying degrees

Although detailed fingerprints were not generated for river dolphins, the profile scores in Fig. 6A provide a window into their evolutionary history. The Indus river dolphin (Platanista), which belongs to one of the earliest-diverging odontocete lineages, retained a strong marine-like profile despite being an obligate freshwater species today — consistent with the marine fossil record of Platanistidae, whose modern representative is now a freshwater specialist. By contrast, the Amazon river dolphin (Inia) and the Yangtze river dolphin (Lipotes) showed lower marine scores but retained high aquatic-dependence scores, consistent with more thorough secondary freshwater adaptation in these lineages.

#### Ancestral branches trace stepwise assembly of marine-like fingerprints

Fig. 6A displays profile scores for selected ancestral branches, which we interpret as rate-based genomic profiles rather than direct habitat reconstructions. Gene-level decompositions of these ancestral profiles (Supplementary Fig. S3) reveal how the marine and aquatic fingerprints were assembled through different gene combinations at successive evolutionary stages.

The cetacean trajectory illustrates this stepwise logic most clearly. The cetacean–hippopotamus ancestor was nearly absent on the marine axis (profile 0.006) but already carried a moderate aquatic-like fingerprint (0.712), with contributions from epithelial, transport, circadian and regulatory genes including TGM3, LRP2, PER1, ATF7IP2 and WDR93. Crown Cetacea strengthened this aquatic-like background (0.877) and gained an intermediate marine-like profile (0.408), adding epithelial, sensory and vascular components such as KRTDAP, OR52W1, SSTR4 and CD34. Deeper cetacean branches then assembled strong marine-like profiles, but through different gene combinations: Mysticeti (0.998) emphasized TGM3, KRTDAP, MYH13, KRT17 and HIGD1B, whereas Odontoceti (0.930) emphasized KRT17, IL36G, TGM3, TPX2 and WDR93. Marine commitment thus arose from an aquatic-like precursor and was elaborated differently in baleen and toothed whale lineages.

Pinniped ancestral branches told a different story. Crown Pinnipedia lacked strong marine-like or aquatic-like fingerprints (marine 0.020; aquatic 0.122), whereas downstream subclades acquired them unevenly. Phocidae assembled both strong marine-like (0.870) and aquatic-like (0.976) profiles, with marine contributions from TGM3, OMP, MYH7B, SDR9C7 and KCNK18. Otarioidea and Otariidae remained weak, but Otariinae later acquired a strong marine-like fingerprint (0.991) through a different gene combination — MYH7B, MYH13, EPOR, CAPN14, OMP and SSTR4 — while retaining a weaker aquatic-like profile (0.339). The pinniped marine signal is therefore not a single ancestral programme inherited from crown Pinnipedia, but a set of unevenly assembled subclade fingerprints.

Sirenian, bear-lineage and otter-lineage ancestral branches all showed weak or partial profiles, despite the marine ecology of some of their extant descendants. This contrast supports the interpretation that the polar bear and sea otter represent marine-edge fingerprints assembled near the tips of their lineages, rather than deeply inherited marine genomic states.

Together, the species and ancestral fingerprints show how a shared slow-rate background is resolved into distinct biological routes: core cetacean and pinniped profiles, heterogeneous pinniped subclades, marine-edge mammals, secondary freshwater cetaceans, and stepwise ancestral shifts — each assembled from different combinations of the same broader gene vocabulary.

## 3. Discussion

Our results show that the dominant genomic signature of mammalian returns to the sea is not a shared adaptive toolkit but widespread evolutionary slowing — a broad directional signal consistent with strengthened functional constraint. This slowing is shared across independently evolved marine lineages, persists after removing cetaceans, and extends to general aquatic dependence. Yet it does not reduce to a fixed set of genes or pathways: compact genomic fingerprints reuse individual genes across lineages but do not preserve recurrent functional modules, and different marine mammals assemble their marine-like profiles through distinct gene combinations. Marine mammal genome evolution is therefore directional but not modular, combining a shared constraint landscape with clade-weighted resolution into biological fingerprints.

This finding reframes how molecular convergence should be read in marine mammals. Previous genome-wide studies have identified parallel amino-acid substitutions and individual candidate genes showing signatures of positive selection or accelerated evolution ^2,4,6^. Our results do not contradict these findings; rather, they place them in a wider genomic context. Individual genes may accelerate — and the fast-side components of the compact fingerprints, concentrated in body-surface, sensory and ion-channel genes, are consistent with remodeling of external interfaces during aquatic life. But the prevailing genome-wide pattern across more than a thousand trait-associated genes is deceleration, not acceleration. Diving, thermoregulation, osmotic balance, oxygen transport and sensory ecology all impose functional requirements that limit how far molecular systems can drift in water. Targeted innovation at selected loci and strengthened constraint across a broad physiological background are complementary layers of the same evolutionary process, not competing explanations.

The dependence of compact marine fingerprints on cetaceans deserves careful interpretation. Cetaceans are the most species-rich, most deeply aquatic and most physiologically committed marine mammals in the dataset. Their disproportionate contribution to the compact fingerprint likely reflects the depth and consistency of their rate-shift profiles — a biological consequence of long and deep aquatic commitment, expressed statistically as more coherent rate profiles. Removing cetaceans weakened compact prediction but left the broad slow-rate signal largely intact, demonstrating that the constraint landscape is not cetacean-specific even though the sharpest fingerprint resolution depends on cetacean sampling. This layered behaviour — broad signal surviving clade removal, compact signal attenuated — may be a general feature of convergent evolution: the underlying directional trend is widely shared, while its compression into small gene sets requires the richest and most deeply adapted contributors.

The species-level and ancestral decompositions add biological resolution to these aggregate patterns. In cetaceans, the cetacean–hippopotamus ancestral branch was nearly absent on the marine axis but already carried a moderate aquatic-like fingerprint, whereas crown Cetacea strengthened this aquatic-like background and gained an intermediate marine-like profile. Strong marine-like fingerprints appeared later in Mysticeti and Odontoceti, but with different gene combinations. This trajectory is consistent with a stepwise ecological transition documented in the fossil record, from early semi-aquatic archaeocetes to fully marine whales ^2,57^, while emphasizing that our ancestral estimates are genomic profile trajectories rather than direct habitat assignments. The moderate aquatic-like signal before a strong marine-like fingerprint raises the possibility of a prolonged freshwater or nearshore stage before full marine commitment — a phase that may be unevenly represented in the fossil record ^2,57^. River dolphins show how present habitat, ancestry and genomic profile can decouple. The Indus river dolphin retained a strong marine-like profile despite being an obligate freshwater species today, consistent with the known marine fossil record of Platanistidae and a relatively recent freshwater colonization ^58^. By contrast, the Amazon river dolphin and the Yangtze river dolphin showed weaker marine-like but strong aquatic-dependence profiles, consistent with more thorough secondary freshwater adaptation. These contrasts show that current habitat alone cannot serve as a simple proxy for genomic history: freshwater cetaceans retain different degrees of marine-like signal depending on lineage history and timing of secondary freshwater transition.

Pinnipeds provide the clearest example of heterogeneous post-crown fingerprint assembly. The crown pinniped branch lacked strong marine-like or aquatic-like fingerprints, whereas strong profiles appeared later and unevenly within the group. Phocidae acquired both marine-like and aquatic-like fingerprints, whereas Otariinae acquired a strong marine-like but weaker aquatic-like profile; Otarioidea and Otariidae remained weaker in the internal-branch projections. This pattern does not prove independent marine origins, but it is consistent with repeated or uneven acquisition of marine-like genomic states after the divergence of major pinniped subclades. Fossils such as Puijila and Potamotherium ^59,60^, together with indirect molecular evidence from taste-receptor pseudogenization ^61^, provide a paleobiological context in which early pinniped relatives occupied freshwater or nearshore settings before the fully marine profiles seen in extant subclades.

Sirenians and marine-edge carnivores illustrate the other end of this spectrum. Sirenian ancestral branches showed weak or partial marine-like fingerprints, consistent with the freshwater and nearshore complexity of early sirenian evolution. The brown bear–polar bear and otter-related internal branches likewise showed weak profiles, whereas the extant polar bear and sea otter carried distinct marine-edge fingerprints built from largely different gene sets than those of cetaceans and pinnipeds. These cases suggest that ecological marine membership can arise through recent or partial shifts near terminal lineages, without producing the deeply aquatic genomic profiles seen in cetaceans and some pinniped subclades — underscoring why the two-layer framework that separates marine specialization from general aquatic dependence is essential.

Several limitations should be kept explicit. Freshwater and semi-aquatic taxa are undersampled relative to marine lineages, limiting power to resolve freshwater-specific adaptations. Rate-based metrics capture shifts in evolutionary constraint but may miss positive selection that does not alter global rates, and they do not directly measure selection coefficients. The functional interpretations of individual predictor genes are hypothesis-generating, informed by existing literature and coefficient direction, but not experimentally validated. Ancestral-branch profiles are rate-based genomic inferences and should not be read as direct habitat assignments, although their alignment with fossil evidence strengthens confidence in the inferred trajectories.

In summary, mammalian returns to the sea were shaped not by a single shared adaptive programme but by a broad directional constraint landscape that different lineages resolved into distinct genomic fingerprints. This pattern suggests that molecular convergence in complex ecological transitions may often be directional before it is modular: repeated environments can impose similar constraints across many genes, while the visible gene-level fingerprints remain clade-weighted, historically contingent and only partly shared. Similar logic may apply to other repeated transitions such as flight, fossoriality and herbivory, where convergence may be written less as a universal toolkit than as a shared direction of genomic change.

## 4. Methods

### 4.1 Species sampling, genome data and trait framework

We started from TOGA-derived coding-gene alignments generated by the Michael Hiller Lab, using the human hg38 assembly as the reference genome ^62^. The nominal coding-gene alignment set contained 17,434 genes. Alignment data were first reduced to one sequence per species where multiple assemblies or subspecies were available and were then subjected to codon-aware quality filtering. Codon columns represented by non-gap codons in fewer than 70% of species were removed as complete three-nucleotide codons to preserve reading frame. For each gene and species, effective sequence length was defined as the number of nucleotide sites represented by A/C/G/T after filtering; sequences with effective length below 10% of the corresponding alignment length were treated as missing.

Species-level sampling was then curated using missing-gene counts, genome quality, phylogenetic placement and ecological representation. Missing-gene counts were used as a quality-control metric and to flag low-completeness genomes, but final species inclusion was manually curated rather than determined by a single automatic threshold. Where multiple subspecies or assemblies represented the same species, we retained the representative with the best data completeness and phylogenetic suitability where possible; where both wild and domesticated forms were available, wild forms were preferred for the comparative analyses. Phylogenetically or ecologically important marine and aquatic taxa could be retained despite higher missingness to preserve coverage of key lineages. This curation yielded the final 302-species dataset used for downstream analyses.

The final dataset comprised 302 mammalian species represented as terminal taxa in the species tree and corresponding branch-coordinate rate matrices. Each species was assigned to one of four curated ecological categories: marine, non-marine aquatic, semi-aquatic or terrestrial/background. The final category counts were 51 marine species, 12 non-marine aquatic species, 13 semi-aquatic species and 226 terrestrial/background species. All 302 species in the final trait table were represented among terminal branches used in the downstream analyses. Full per-species trait assignments and curated marine-source attributions are listed in Supplementary Table S1.

Marine membership was treated as a curated binary trait, not as a threshold on aquaticity score. Marine species were assigned using established marine mammal authorities, primarily Berta et al. (2015) ^1^ and the Society for Marine Mammalogy Committee on Taxonomy (2026) ^63^. This marine category included fully aquatic marine lineages as well as marine-associated taxa with lower aquaticity scores, such as the polar bear and sea otter, while retaining marine identity as a taxonomic/ecological category separate from the quantitative aquaticity score. For the marine-specialization analyses, all marine species were coded as trait-positive, and all non-marine species were coded as background.

General aquatic dependence was curated independently from marine membership using the five-dimensional scoring framework described below. Non-marine aquatic taxa were non-marine species assigned to the strongly aquatic state in the primary aquatic-dependence endpoint, whereas semi-aquatic taxa were non-marine species assigned to an intermediate aquatic-dependence state. The strongly aquatic positive set therefore included marine and non-marine aquatic taxa. Semi-aquatic labels were excluded from the primary binary aquatic-dependence endpoint to preserve an unambiguous two-state contrast. Thus, the primary binary aquatic-dependence endpoint contrasted 63 strongly aquatic species against 226 terrestrial/background species, with 13 semi-aquatic species excluded. Alternative codings that treated semi-aquatic taxa as aquatic-positive or as terrestrial/background were used only as coding-sensitivity analyses.

### 4.2 Five-dimensional aquaticity scoring and PCA validation

General aquatic dependence was quantified using a five-dimensional aquaticity scoring framework. The five dimensions were foraging medium, locomotion or escape medium, reproduction or nursery dependence, morpho-physiological specialization, and aquatic time budget. For foraging medium, locomotion or escape, and reproduction or nursery, ordinal scores ranged from 0 to 3, corresponding broadly to no meaningful aquatic dependence, weak or occasional water association, substantial but non-exclusive aquatic dependence, and strong or predominant aquatic dependence. Morpho-physiological specialization was scored from 0 to 5 to capture anatomical and physiological specialization for aquatic life: 0 indicated no clear aquatic specialization, intermediate scores indicated increasing aquatic-related specialization, score 4 denoted paddle-like limbs or comparably advanced aquatic structural adaptation, and score 5 denoted extreme obligate aquatic specialization, including loss of hind limbs. Aquatic time budget was scored from 0 to 4, ranging from negligible time spent in water to score 4 for species spending virtually their whole life in water. The summed aquaticity score therefore ranged from 0 to 18. Higher values indicate stronger dependence on aquatic environments, but the summed score was not used to define marine membership. Per-species dimension scores, summed totals and scoring criteria are listed in Supplementary Table S1.

The summed aquaticity score was used to organize the gradient of general aquatic dependence. Of the 302 species, 226 scored zero. No sampled species scored 5 or 6, producing an empirical gap that was used to operationalize the semi-aquatic/aquatic boundary for the primary binary aquatic-dependence endpoint. Species scoring above this gap (score ≥ 7) coincided with the curated strongly aquatic set, comprising marine and non-marine aquatic taxa, used as aquatic-positive for the primary binary aquatic-dependence endpoint; terrestrial/background species were coded as negative and semi-aquatic species were excluded. Marine species occupied much of the high-score range, including all 8 species at score 15 and all 37 species at score 18; however, marine identity was not defined by score alone. For example, the polar bear and sea otter were retained as marine species despite lower aquaticity scores of 7 and 13, respectively.

To validate whether the five scoring dimensions behaved as a single dominant aquaticity gradient, we performed principal component analysis on the five raw dimension scores across the 302 species. The dimension scores were centered and scaled before PCA using prcomp(…, center = TRUE, scale. = TRUE). The sign of PC1 was oriented so that larger PC1 values corresponded to higher summed aquaticity scores. PC1 explained 97.0% of total variance and PC2 explained 2.2%. PC1 was almost perfectly correlated with the summed aquaticity score (r = 0.99992; R² = 0.99984), supporting use of the simple summed score as a compact quantitative axis of general aquatic dependence, rather than redefining aquaticity by PCA-derived weights.

### 4.3 Species tree, branch-coordinate matrices, GBI construction and deterministic branch-state labels

The species tree for the final 302-species dataset was inferred from single-copy coding genes in the curated alignment set. Genes with alignment length greater than 2,000 bp were used for this step, yielding 2,275 candidate genes for species-tree inference. Maximum-likelihood gene trees were inferred with IQ-TREE2 (v2.1.3) using automatic nucleotide-substitution model selection, 1,000 bootstrap replicates, and a partition scheme distinguishing first-plus-second codon positions from third codon positions ^64^. The species tree was then inferred from the resulting gene trees using the coalescent method implemented in ASTRAL III ^65^.

Gene-wise branch lengths were estimated by fixing each gene-tree topology to the final 302-species tree and estimating branch lengths from the corresponding coding-sequence alignment using BaseML from the PAML package ^66^. Branch lengths were estimated under a REV/GTR-like nucleotide substitution model with discrete gamma rate heterogeneity ^67–69^. Because different genes lack different species, the reduced taxon set available for a gene can cause adjacent species-tree branches to collapse into fused gene-tree branches. We therefore used SplitAligner to project gene-wise branch lengths onto the common species-tree branch-coordinate system ^8^. Branch lengths were retained only when a gene-tree branch could be assigned uniquely to a single species-tree branch coordinate; fused or non-uniquely assignable branches were treated as missing for the corresponding individual species-tree branch coordinates rather than being assigned to adjacent branches.

Gene-by-branch evolutionary-rate features were derived from the resulting branch-coordinate-aligned branch-length matrices produced by SplitAligner ^8^. The endpoint-fix branch-coordinate matrix used for this study comprised 17,432 genes across 601 branches of the 302-species mammalian tree.

Gene–branch interaction (GBI) values were computed to express each gene’s branch length relative to gene-level and branch-level background rates, so that a GBI value reflects a gene-specific deviation from the genome-wide rate of a branch rather than an absolute branch length ^9^. Within each gene, branch lengths at or above the 97.5th within-gene percentile were set to missing before summary statistics were computed, limiting the influence of a small number of extreme per-gene values. Branch effects were computed as branch-wise means over non-missing entries of the trimmed branch-length matrix, and gene effects were computed as gene-wise means over non-missing entries of the same trimmed matrix ^70^. The final GBI value for gene *g* on branch *b* was computed as

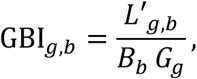

where *L*′_g,*b*_ is the trimmed branch length, *Bb* is the branch effect, computed as the mean trimmed branch length for branch *b* across genes, and *G_g_* is the gene effect, computed as the mean trimmed branch length for gene *g* across branches. Non-finite values generated by missing branch lengths, missing effects or division by zero were retained as missing. GBI construction is trait-independent: it uses branch-coordinate branch lengths and does not use trait labels at any step.

Deterministic ancestral-state branch-state labels were assigned separately for each trait by maximum-parsimony ancestral-state reconstruction on the species tree using ‘`castor::asr_max_parsimony`’ ^71,72^. Ties in the ancestral-state support matrix were resolved by selecting the first tied state in the fixed state ordering used for that trait (‘`ties.method = “first”`’), making the reconstruction fully deterministic. Repeated reconstruction confirmed this determinism: identical branch-state vectors were recovered across 20 repeated reconstructions for every trait. Node states were mapped to branches using the reconstructed state of the descendant node of each branch, assigning one trait-state label to each of the 601 species-tree branches. For the marine trait, this produced 97 marine-state branches and 504 background-state branches. For the binary aquatic-dependence endpoint, deterministic reconstruction produced 114 aquatic-state branches, 474 background-state branches and 13 intermediate semi-aquatic branches; the 13 intermediate branches were excluded before binary branch-level contrasts.

These branch-state labels are trait-derived but deterministic and frozen. They were not estimated from gene-level rate data, and they were computed once and held fixed. Throughout all single-gene screens, cross-validation folds and sensitivity analyses reported below, the GBI matrix was treated as a frozen trait-independent feature-construction layer, whereas deterministic branch-state labels were treated as frozen trait-derived annotation layers. The nested procedures described in Sections 4.6 and 4.8 re-derived supervised candidate-gene selection within folds but did not recompute the GBI matrix or the deterministic branch-state labels.

### 4.4 Global branch-level single-gene screening and positive-count-matched permutation control

The global branch-level single-gene screens reported in Fig. 3 and Fig. S2 are association and directional screens applied to the GBI matrix; they are not LASSO models and are not cross-validated predictive analyses. For each gene, GBI values were compared between deterministic trait-state and background-state branches (Section 4.3) using a Welch two-sample t-test ^73^. Missing GBI values were removed on a gene-by-gene basis before each test, and a gene was tested only when more than one finite value remained in each branch-state group. These single-gene screens therefore used available non-missing GBI values directly and did not use the terminal-mean imputation applied to LASSO input matrices (Section 4.7).

For each Welch test, nominal *P* values were calculated from the two-sided Welch test, whereas the slow/fast direction was determined from the signed background-minus-trait mean contrast,

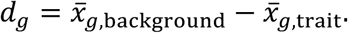

Here *x̄_g_*_,background_ and *x̄_g_*_,trait_ denote the mean finite GBI values of gene *g* on deterministic background-state and trait-state branches, respectively. Under this convention, a positive *d_g_* corresponds to lower mean GBI on trait-state branches than on background branches and is reported as a slow-direction gene; a negative *d_g_* corresponds to higher mean GBI on trait-state branches and is reported as a fast-direction gene. This is the slow/fast convention used throughout Fig. 3 and Fig. S2. For signed screening plots, the Welch test statistic was displayed with its sign assigned by the background-minus-trait contrast *d_g_*.

False-discovery-rate control ^74^ was applied within each screen using a sorted-*P* Benjamini–Hochberg selection rule. Let *p*_(1)_ ≤ *p*_(2)_ ≤ ⋯ ≤ *p*_(*m*)_ denote the sorted nominal *P* values for *m* tested genes. We identified the largest rank *k* satisfying

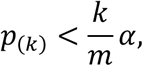

with *α* = 0.01, and genes with rank ≤ *k* were declared FDR-significant. If no rank satisfied this criterion, no genes were declared significant in that screen. The marine-binary screen used all 601 species-tree branches and tested 17,258 genes. The primary binary aquatic-dependence screen excluded the 13 branches assigned the intermediate semi-aquatic state and used the remaining 588 branches to test 17,298 genes. Numerical summaries of significant slow-and fast-direction genes are reported in Fig. 3, Fig. S2, and Supplementary Tables S2 and S4. Sensitivity screens used the same Welch t-test, signed background-minus-trait direction convention and FDR threshold.

A positive-count-matched permutation control was applied to the endpoint-fix drop-cetacean marine single-gene screen to test whether the observed FDR-significant-gene count and slow-direction proportion among significant genes could be explained by the reduced number of remaining marine-positive terminal labels after cetacean removal (Fig. 5B). The matched terminal-label design comprised 17 marine-positive terminal species and 251 background terminal species. Here, “positive-count-matched” means that each permutation preserved the observed number of positive terminal labels; it does not imply exhaustive enumeration of all possible label assignments.

For each permutation, 17 eligible terminal species were sampled without replacement as the positive set, and the remaining 251 eligible terminal species were assigned as background. For the permutation control, the deterministic ancestral-state reconstruction procedure described in Section 4.3 was rerun on the permuted terminal labels. The resulting node states were mapped to branches, run-specific excluded branch states were omitted, and the branch-level Welch t-test and sorted-*P* FDR rule with *α* = 0.01 were applied as in the observed screen. Each permutation therefore preserved the observed positive-count terminal-label design while randomizing which terminal species carried the positive label, with downstream branch-state reconstruction and branch-level screening recomputed under that permuted labeling.

Two statistics were recorded for each permutation: the number of FDR-significant genes and, when at least one FDR-significant gene was selected, the proportion of significant genes with positive signed background-minus-trait contrasts, corresponding to the slow direction. A total of *n*perm = 200 permutations was run. The slow-direction proportion was treated as undefined for permutations in which no FDR-significant gene was selected.

Empirical *P* values were calculated as upper-tail one-sided probabilities with a +1 correction. For the significant-gene count,

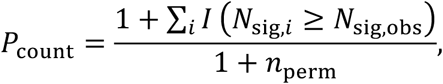

where *N*_sig,obs_ is the observed number of FDR-significant genes and *N*_sig,*i*_ is the corresponding number in permutation *i*. For the slow-direction proportion,

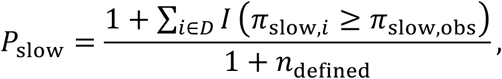

where *D* is the set of permutations with at least one FDR-significant gene, *π*_slow,obs_ is the observed slow-direction proportion, *π*_slow,*i*_ is the corresponding proportion in defined permutation *i*, and *n*_defined_ is the number of permutations with at least one FDR-significant gene and therefore a defined slow-direction proportion. Thus, *P*_slow_ was conditional on permutations with a defined slow-direction proportion. Observed and null summaries of the drop-cetacean marine screen are reported in Fig. 5B and Supplementary Table S6. Branch-level positive/background counts from the per-permutation ancestral-state reconstructions were retained as provenance and QC values; the public matched design for Fig. 5B is the terminal-label design of 17 positives and 251 negatives.

### 4.5 Two-stage LASSO framework and evidence levels

Trait-associated genome-wide rate structure was modelled in two stages. In Stage 1, candidate genes were identified by the branch-level single-gene screen described above: genes whose GBI values differed between trait-state and background-state branches at a Benjamini–Hochberg false-discovery-rate threshold of FDR ≤ 0.01 ^74^. In Stage 2, the GBI values of the Stage-1 candidate genes were used as features in penalized logistic regression models fitted with LASSO on terminal species ^75,76^.

Two trait endpoints were modelled. The marine model predicted marine specialization, with marine species coded as trait-positive and all non-marine species coded as background. The binary aquatic-dependence model predicted strong general aquatic dependence, with strongly aquatic species coded as trait-positive and terrestrial/background species coded as negative; semi-aquatic species were excluded from the primary binary aquatic-dependence LASSO sample domain. Alternative aquatic codings that treated semi-aquatic taxa as aquatic-positive or as background were analysed only as coding-sensitivity comparisons and were not used for the primary aquatic-dependence model.

We kept five evidence layers distinct. First, the global branch-level single-gene screens in Fig. 3 and Fig. S2 tested each gene independently against deterministic branch-state labels using the GBI matrix and branch-level contrasts described above. These screens define full-data genome-wide association patterns and, where used for full-data architecture summaries, full-data candidate-gene sets; they are not LASSO validation analyses. Second, predictive performance in Fig. 2 and Fig. 5A was estimated only by nested supervised feature-selection gLOOCV, in which supervised candidate-gene selection was repeated inside each outer cross-validation fold. Third, selected-predictor architecture in Fig. 4 and Supplementary Table S5 was summarized by final full-data LASSO fits performed after validation; these full-data fits describe selected genes and coefficients but are not cross-validated performance estimates. Fourth, the positive-count-matched permutation control in Fig. 5B evaluates the branch-level single-gene screen and is not a LASSO validation analysis. Fifth, predictor-turnover analyses in Fig. 5C and Supplementary Table S6 compare corrected full-data predictor sets against comparison-specific candidate-gene nulls. Throughout these analyses, the GBI matrix and deterministic branch-state labels were held fixed as described above; only supervised candidate-gene selection and LASSO model fitting were re-estimated within validation folds.

### 4.6 Genus-level leave-one-out cross-validation and sensitivity-run definitions

Predictive performance was estimated by genus-level leave-one-out cross-validation over terminal species. In each outer fold, all eligible terminal species from one genus were held out as the test set, and all remaining eligible terminal species formed the terminal training set. Holding out entire genera, rather than individual species, reduced the chance that close relatives of held-out species remained in the training set. For the marine model, all 302 terminal species were eligible. For the primary binary aquatic-dependence model, the 13 semi-aquatic taxa were excluded before training, held-out prediction, preprocessing and AUC calculation, leaving 289 evaluable terminal species. Folds in which the held-out genus contained no eligible terminal species after run-specific exclusion rules were applied did not contribute finite out-of-fold predictions or AUC calculations.

Within each outer fold, supervised branch-level t-test/FDR candidate-gene selection was repeated using the fold-specific branch-level screening table described below. The GBI matrix was fixed as a trait-independent rate-feature layer, whereas deterministic branch-state labels were fixed as trait-derived annotation layers. Candidate genes with FDR ≤ 0.01 in the training fold were passed to terminal-only LASSO fitting for that fold.

Discrimination was summarized from pooled finite out-of-fold predictions using a rank-based Mann–Whitney/Wilcoxon-equivalent AUC statistic with average ranks for ties ^77^. Higher predicted probabilities were interpreted as stronger support for the run-specific positive class. Numerical performance summaries, including nested AUC values, fold-specific candidate-gene counts and selected-predictor counts, are reported in Fig. 2, Fig. 5A and Supplementary Table S6.

The Fig. 5A sensitivity analyses used the same genus-level cross-validation, nested supervised feature-selection and terminal-only LASSO procedure. Marine-specialization runs were defined as follows: marine baseline, 51 positives and 251 negatives; drop cetaceans, 17 positives and 251 negatives; Cetacea only, 34 positives and 268 negatives; and Pinnipedia only, 12 positives and 290 negatives. In drop-clade analyses, the removed clade was excluded from both training and held-out evaluation. In clade-only analyses, the focal clade was coded as the run-specific positive class and all other eligible terminal species were coded as background; AUCs for these runs were computed against run-specific labels rather than the global marine label.

Binary aquatic-dependence runs were defined as follows: aquatic baseline, 63 positives and 226 negatives; no Cetacea, 29 positives and 226 negatives; no Pinnipedia, 51 positives and 226 negatives; and no marine-edge taxa, 58 positives and 226 negatives. Marine-edge taxa were defined as *Dugong dugon*, *Enhydra lutris kenyoni*, *Hydrodamalis gigas*, *Trichechus manatus latirostris* and *Ursus maritimus*, comprising three sirenians, the sea otter and the polar bear. In the no-marine-edge binary aquatic-dependence sensitivity run, these taxa were recoded from aquatic-positive to the excluded endpoint category before training, held-out prediction, preprocessing and AUC evaluation.

### 4.7 Missing-value imputation, fold-wise scaling and model fitting for LASSO

LASSO models require a complete numeric design matrix, whereas GBI matrices contain missing values arising from the 97.5^th^-percentile trimming described above and from genes absent on particular branches. For LASSO input only, missing values in the selected-feature matrix were replaced by gene-wise means computed over eligible terminal species; in cross-validation, these means were computed from training terminal species alone. This imputation supplies numeric placeholders for `glmnet` input and is not interpreted as a branch-rate estimate. Unlike the global branch-level single-gene screens described in Section 4.4, which used available non-missing GBI values directly, the LASSO models required numeric placeholders for missing values and therefore used gene-wise mean imputation within the eligible terminal-species set.

All LASSO models were fitted on terminal species only; internal branches were not used as LASSO observations. Features were scaled manually within each fold using means and standard deviations learned from training terminal species alone, and `glmnet` was therefore called with `standardize = FALSE` to prevent additional internal scaling. Features with no observed training-terminal values or zero training-terminal variance in a fold were excluded from that fold before model fitting. Penalized logistic models were fitted with `glmnet` using `family = “binomial”`. The `alpha` argument was not overridden; the `glmnet` default `alpha = 1` was therefore used, corresponding to a pure LASSO penalty. The regularization strength was selected as `lambda.min` from `cv.glmnet` using `type.measure = “deviance”`; inner fold identifiers were defined only over the training terminal species, with one training terminal species per inner fold. In every cross-validation fold, the held-out genus contributed to none of imputation, scaling, lambda selection or model fitting. For the binary aquatic-dependence models, semi-aquatic species with endpoint value 0.5 were excluded from all LASSO sample domains, including training, held-out prediction, imputation, scaling, lambda selection and AUC evaluation.

For final full-data architecture fits, the same imputation, scaling and lambda-selection procedures were applied using all eligible terminal species in the corresponding model. These full-data fits were used only to summarize selected-predictor architecture and coefficient structure, not to estimate held-out predictive performance.

### 4.8 Primary nested supervised feature-selection gLOOCV

Because Stage-1 candidate genes were selected using trait labels, selecting them once on all species and reusing them across cross-validation folds would allow held-out information to influence feature selection and could inflate apparent predictive performance. To prevent this, supervised feature selection was nested inside the outer genus-level cross-validation. For the main Fig. 2 validation and the Fig. 5A sensitivity analyses, branch-level t-test/FDR candidate-gene selection was repeated inside each outer genus-level leave-one-out fold. In each fold, the run-specific inclusion and exclusion rules were first applied, and terminal branches belonging to the held-out genus were removed from the branch-level screening table. Candidate genes passing FDR ≤ 0.01 in this fold-specific branch-level screen were then passed to terminal-only LASSO fitting with fold-wise training-terminal-only imputation, scaling, lambda selection and model fitting.

The same two-sided Welch test and Benjamini–Hochberg FDR ≤ 0.01 candidate-gene rule used in the global branch-level screen were applied within each training fold. Slow/fast direction was assigned using the same signed background-minus-trait mean-contrast convention, so that positive contrasts corresponded to lower mean GBI on trait-state branches than on background branches, matching the slow-direction convention used in Fig. 3.

This nesting was applied to supervised trait-labelled feature selection, not to global phylogenomic feature construction. The GBI matrix was kept fixed as a trait-independent rate-feature matrix, whereas deterministic branch-state labels were kept fixed as trait-derived annotation layers. Internal branches were retained as branch-level screening observations within each fold; only the terminal branches of the held-out genus were removed from the supervised screen. These analyses are therefore nested with respect to supervised feature selection and terminal LASSO fitting, but they are not fully nested end-to-end phylogenetic reconstructions.

As a sensitivity check for the fixed branch-state annotation layer, we repeated the baseline marine and binary aquatic-dependence nested gLOOCV after masking held-out-genus terminal labels before ASR inside each outer fold. The main deterministic branch-state labels were generated with `castor::asr_max_parsimony` as described in Section 4.3. For this fold-wise sensitivity check, held-out-genus terminal labels were set to NA for ASR only, and fold-specific deterministic parsimony reconstruction was performed with `castor::hsp_max_parsimony`, which permits unknown tip states. A no-missing validation confirmed that this `hsp_max_parsimony` implementation exactly reproduced the frozen `asr_max_parsimony` branch-state vectors before label masking. The resulting fold-specific branch-state labels were then used for the same descendant-node branch mapping, branch-level t-test/FDR feature selection and terminal-only LASSO fitting used in the main nested workflow. Baseline AUCs were unchanged for the marine model and only minimally changed for the binary aquatic-dependence model (marine: frozen-ASR 0.936, fold-wise ASR 0.936; binary aquatic-dependence: frozen-ASR 0.826, fold-wise ASR 0.823), indicating that the fixed ASR annotation did not materially affect the baseline validation results.

Because the fold-specific screen re-selects genes from training data, the candidate-gene set differs across folds; fold-level candidate-gene and selected-predictor summaries are reported in Supplementary Table S6. As a consistency check, recomputing the global Stage-1 screen without holding out any genus returned the archived Stage-1 candidate set exactly. Both the marine and binary aquatic-dependence models, and all sensitivity runs described above, used this nested supervised feature-selection procedure, so that the AUC values reported in Fig. 2 and Fig. 5A are out-of-sample with respect to supervised feature selection and LASSO model fitting.

### 4.9 Full-data LASSO architecture and predictor annotation

Final full-data LASSO fits were performed after validation to summarize the architecture of the corrected sparse predictor solutions. These full-data fits used the corresponding full-data Stage-1 candidate genes from the branch-level t-test/FDR screens described in Section 4.4: 1,559 marine-binary candidate genes and 1,227 binary aquatic-dependence candidate genes. The same LASSO implementation framework described in Section 4.7 was then applied: terminal species only, terminal-mean imputation over eligible terminal species, manual scaling computed over the same eligible terminal species, lambda.min selection via cv.glmnet with `family = “binomial”` and `type.measure = “deviance”`, and glmnet fitting with `standardize = FALSE`. For the binary aquatic-dependence model, terminal species with the intermediate endpoint value 0.5 were excluded from the eligible terminal set throughout, as in Section 4.7. The marine full-data fit therefore used 302 eligible terminal species (51 trait-positive and 251 background), and the binary aquatic-dependence full-data fit used 289 eligible terminal species (63 trait-positive and 226 background). These full-data fits are architecture summaries, not cross-validated performance estimates; predictive performance was estimated by the nested gLOOCV procedures described in Sections 4.6 and 4.8 and reported in Fig. 2 and Fig. 5A.

The corrected full-data baseline marine model selected 71 predictors and the corrected full-data baseline binary aquatic-dependence model selected 101 predictors. Of these, 24 predictors were shared between the two final full-data baseline models, 47 were selected only by the marine model and 77 only by the binary aquatic-dependence model, yielding a union of 148 unique predictors (Fig. 4A). A “shared” classification here indicates that the same gene was retained as a non-zero predictor by both final full-data baseline models; it does not imply the same coefficient sign, the same effect size or the same biological role. LASSO coefficients were fitted on the scaled GBI input features described above. For display in Fig. 4B and Fig. 4C, positive model coefficients are labelled as fast-direction coefficients and negative model coefficients as slow-direction coefficients because LASSO coefficients use the direct scaled-GBI feature orientation. This coefficient orientation differs from the signed background-minus-trait contrast used to orient single-gene screening statistics in Section 4.4. The visual emphasis threshold was |coefficient| > 0.1. Coefficient sign and magnitude in this section are model-architecture quantities and should not be interpreted as standalone single-gene branch-rate evidence; the corresponding directional evidence at the single-gene level is described in Section 4.4 (Fig. 3 and Fig. S2).

All 148 corrected full-data predictors are reported in Supplementary Table S5, together with predictor class, marine and binary aquatic-dependence coefficients, coefficient direction, compressed display module, recommended submodule or function, annotation confidence, evidence or source notes, and Fig. 4C display status. The Fig. 4C module-annotated display uses 12 compressed descriptive display modules: Cilia / flagella / reproductive-cell function; Cytoskeletal / muscle / ECM / adhesion; DNA repair / chromatin / cell-cycle; Developmental / transcriptional signaling regulators; Endocrine / circadian / systemic regulation; Epithelial / body-surface interface; Immune / inflammatory regulation; Ion channels / mechanosensation / membrane signaling; Metabolism / redox / lipid handling; Sensory systems; Transport / endocytic / epithelial solute handling; and Vascular / hematologic regulation. Across these 12 main display modules, 118 of the 148 predictors were assigned to a module and are counted in the Fig. 4C circle sizes. Each circle therefore represents the total number of module-assigned predictors in a module-by-partition cell, not only those genes whose names are printed; gene labels in Fig. 4C are representative high-or medium-confidence genes selected for readability. The remaining 30 predictors, including Table S5-only records, low-confidence annotations and unassigned predictors, are reported in Supplementary Table S5 but are not included in the 12-module Fig. 4C circle-size counts. The Fig. 4C module grouping is a descriptive organization of the corrected full-data predictors; it is not a pathway-enrichment test and is not evidence of module-level evolutionary recurrence. Functional enrichment of the genome-wide single-gene screens is described in Section 4.11 (Fig. 3C and Fig. S2; Supplementary Table S3), and module-level turnover relative to candidate-gene null expectations is evaluated separately in Section 4.10 (Fig. 5C).

### 4.10 Predictor turnover and comparison-specific candidate-gene null models

To evaluate whether sparse predictor overlap exceeded expectations from the available candidate-gene background, we compared corrected full-data predictor sets from the corresponding LASSO models. The two main predictor-versus-predictor comparisons shown in Fig. 5C were the marine baseline versus cetacean-only models and the binary aquatic-dependence baseline versus no-Cetacea models. A separate cross-layer comparison between marine baseline LASSO predictors and the drop-cetacean marine-slow single-gene set was retained only as supplementary context and was not treated as the same evidence class as the main predictor-versus-predictor comparisons.

Predictor turnover was quantified at both gene and module levels. Gene-level overlap was measured using the Jaccard index between the two compared predictor sets. Module-level similarity was evaluated using the compressed display-module framework defined for Fig. 4C and Supplementary Table S5. We used two module-level metrics: a module-presence Jaccard index, based on the set of modules represented in each predictor set, and a module-count cosine similarity, based on the vector of predictor counts across modules. Genes without curated module assignments were included in gene-level overlap calculations but were not used in module-level similarity calculations.

Null expectations were generated separately for each comparison. For a comparison between models *A* and *B*, let *C_A_* and *C_B_* denote the candidate-gene sets available to the two corresponding full-data LASSO models before final predictor selection. The gene-level null sampling universe was defined as

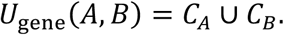

Random gene sets matching the observed predictor-set sizes of models *A* and *B* were sampled without replacement from this comparison-specific candidate-gene union, and the resulting gene-overlap Jaccard distribution was used as the gene-level null expectation. For module-level turnover, null sampling was restricted to the module-annotated subset of the same comparison-specific candidate-gene union,

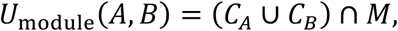

where *M* denotes genes with curated module annotations. Module-level random sets were likewise sampled without replacement and matched the numbers of module-annotated genes in the two observed predictor sets, rather than the total predictor-set sizes. This ensured that module-level null expectations were evaluated against the annotation-eligible candidate-gene background available to that specific comparison, rather than against the full genome, all annotated genes, or the observed selected predictors alone.

For each comparison, 10,000 random permutations were used to generate null distributions for gene-overlap Jaccard, module-presence Jaccard and module-count cosine similarity under matched-size random sampling. We report the observed value for each metric together with the null median, 95% null interval and empirical *P* value in Fig. 5C and Supplementary Table S6. The 95% null interval was defined as the 2.5th and 97.5th percentiles of the permutation distribution. Empirical *P* values were calculated as upper-tail one-sided probabilities with a +1 correction,

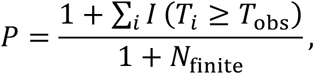

where *T*_obs_ is the observed overlap or similarity statistic, *T_i_* is the corresponding statistic from permutation (i), *N*_finite_ is the number of finite null statistics, and *I*(.) is the indicator function. This tested whether the observed overlap or similarity exceeded the comparison-specific null expectation. All reported null summaries had 10,000 finite permutation values. These empirical *P* values were used as comparison-wise null diagnostics for the turnover metrics rather than as a genome-wide multiple-testing screen.

For the supplementary cross-layer comparison, the observed sets were the corrected full-data marine baseline LASSO predictors and the drop-cetacean marine-slow FDR genes. Random gene sets were sampled from the union of the marine baseline FDR candidate genes and all FDR-significant genes from the drop-cetacean marine screen, including both slow-and fast-direction significant genes; this universe was not the full set of tested genes and was not restricted to the observed slow-direction genes alone. Module-level sampling for this cross-layer comparison was again restricted to genes with curated module annotations in the same union. Because this comparison contrasts a sparse LASSO predictor set with a single-gene slow-direction set, it was not included in the main Fig. 5C panel and is reported only as supplementary context.

These turnover analyses were performed on corrected full-data predictor sets and curated module annotations after model validation. They were therefore used to interpret predictor-set overlap relative to candidate-gene background expectations, not to estimate cross-validated predictive performance.

### 4.11 Functional enrichment analysis

Functional enrichment was used as descriptive annotation of the single-gene screening results. Inputs were the four directional FDR-significant gene sets from the branch-level t-test screens described in Section 4.4: marine-slow (n = 1,366), marine-fast (n = 193), aquatic-slow (n = 1,055) and aquatic-fast (n = 172).

Enrichment results were archived direct exports from the STRING web tool, using the Homo sapiens network in STRING v12.0 (exported 20 May 2026) ^78^. Gene symbols were submitted against the Homo sapiens network, and archived species-9606 protein identifiers confirm the organism setting. For each directional gene set, the archived export records genes mapped to STRING species-9606 protein identifiers; unmapped or unrecognized symbols, if any, were not represented in the STRING enrichment exports. No custom local background was supplied; enrichment used STRING’s default organism/network background, and term-specific background counts are retained from the exported tables. The archived exports preserve STRING-provided FDR values, observed gene counts and background gene counts per term. Because only direct STRING web exports were archived, we report STRING-provided FDR values as exported and do not re-label the underlying STRING procedure as a local Fisher or hypergeometric analysis. Because enrichment was performed using STRING’s default background rather than an analysis-specific testable-gene background, enrichment results were used for descriptive annotation rather than formal comparisons of functional coherence between gene sets.

Supplementary Table S3. contains the full archived STRING outputs for all four directional sets across STRING-exported categories, including Monarch, Gene Ontology, Reactome, KEGG, InterPro, TISSUES, UniProt Keywords and other STRING-derived categories. Terms were considered enriched according to the STRING-provided threshold of FDR ≤ 0.05. Fig. 3C displays a curated visualization subset of enriched marine-slow STRING/Monarch phenotype and measurement terms, selected by prioritizing hematological, erythrocyte, hemoglobin, platelet, blood-cell, cardiovascular, anthropometric/body-measurement and protein-measurement terms while omitting redundant or less informative terms. Each displayed term is plotted by observed gene count and −log_1B_(STRING FDR). The full exported enrichment outputs are reported in Supplementary Table S3.

Aquatic-slow enrichment was summarized in the same phenotype/measurement annotation space and using the same STRING-provided FDR threshold. Among STRING/Monarch phenotype and measurement terms passing this threshold in the aquatic-slow output, one broad term, Hematological measurement (EFO:0004503), was retained for descriptive summary and is reported in Fig. S2 and Supplementary Table S3. No size-matched marine-versus-aquatic enrichment-coherence test was run, so the difference between the broader marine-slow display and the narrower aquatic-slow result is reported descriptively only and is not interpreted as differential functional coherence.

This section provides descriptive annotation of genome-wide screening-derived gene sets; it is not mechanistic proof, not the Fig. 4C predictor-module annotation, and not evidence of module-level recurrence, which is evaluated by the candidate-gene null turnover analyses described in Section 4.10.

### Use of large language models and AI-assisted tools

No large language model or AI-assisted tool is listed as an author. Large language models and AI-assisted tools, including OpenAI ChatGPT models, OpenAI Codex and Anthropic Claude models, were used as auxiliary aids for manuscript drafting, wording refinement, code drafting and review under author-written specifications, figure and table caption review, and inspection of author-curated quality-control packages. No large language model or AI-assisted tool was used as primary data, independent scientific evidence, or an autonomous source of statistical computation, model fitting, P-value calculation, permutation, ancestral-state reconstruction or null-model generation. All analyses were executed by author-written or author-supervised scripts and checked against archived scripts, tables and source files. AI-assisted outputs were reviewed, edited and, where necessary, corrected by the authors before inclusion. The authors made all final decisions and take full responsibility for the study.

## Author contributions

J.W. and T.Y. conceived the study and jointly developed the trait framework. J.W. developed the analytical framework, designed and performed the computational analyses, interpreted the results, prepared the figures, and wrote the manuscript. T.Y. contributed to trait design, species curation, aquaticity-score calibration, and biological interpretation. N.K. contributed marine mammal ecological and paleobiological expertise to the aquaticity scoring framework and species-level trait assignments. H.K. contributed to statistical framing, including the choice of the t-test screening framework, and to discussion of GBI scaling and evolutionary-rate interpretation. All authors reviewed and approved the final manuscript.

## Materials & correspondence

Correspondence and requests for materials should be addressed to J.W., N.K. or H.K.

## Data availability

Study-generated data supporting this manuscript have been deposited in Dryad under DOI https://doi.org/10.5061/dryad.dz08kpsd4. The dataset is currently under embargo and will be made publicly available upon publication. The deposited dataset includes curated trait tables, aquaticity scores, species-tree files, fixed-topology gene trees, fixed-topology branch-length outputs, SplitAligner branch-coordinate matrices, gene–branch interaction matrices, deterministic branch-state labels, single-gene screening outputs, LASSO outputs, permutation-control outputs, predictor-turnover null outputs, functional-enrichment exports, Source Data files and Supplementary Tables. External TOGA-derived coding-gene alignments from the Michael Hiller Lab were used as starting genomic resources but are not redistributed in the Dryad dataset; source information and processing manifests are provided instead.

## Code availability

Custom scripts used for data processing, branch-coordinate matrix construction, GBI calculation, deterministic branch-state labelling, single-gene screening, nested gLOOCV LASSO analyses, fold-wise ASR sensitivity checks, positive-count-matched permutation controls, predictor-turnover null analyses, functional-enrichment processing, figure generation and package validation are publicly available at https://github.com/wujiaqi06/marine_mammal under release tag v1.1-analysis-update; the frozen endpoint-fix baseline remains available as v1.0-endpoint-fix. Large input and output data required by these scripts have been deposited in Dryad at https://doi.org/10.5061/dryad.dz08kpsd4. SplitAligner is available separately at https://github.com/wujiaqi06/SplitAligner.

## Ethics statement

This study used publicly available or previously generated genomic resources and did not involve new animal experiments, human participants, field sampling, or collection of new biological specimens. No ethics approval was required.

## Supplementary figures and tables

**Supplementary Fig. S1.**
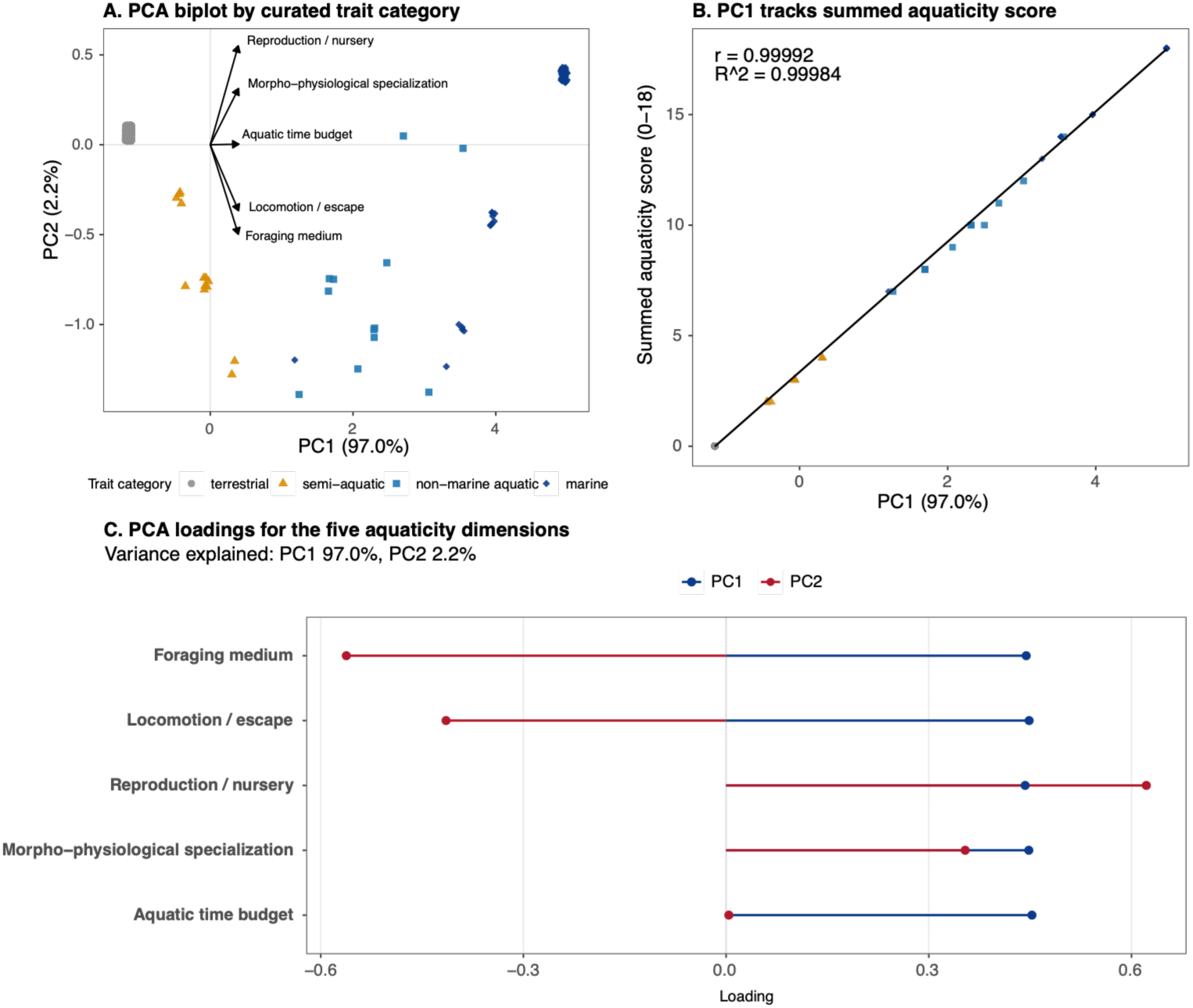
The summed aquaticity score captures a single dominant gradient of general aquatic dependence. (A) PCA biplot of the five aquaticity scoring dimensions, with species coloured by curated ecological category: terrestrial (grey), semi-aquatic (orange), non-marine aquatic (blue squares) and marine (blue diamonds). The large grey cluster at the origin represents the 226 terrestrial species that scored zero on all dimensions. (B) PC1 tracked the summed aquaticity score almost perfectly (r = 0.99992; R² = 0.99984), supporting use of the simple summed score rather than PCA-derived weights as the quantitative axis of general aquatic dependence. (C) PCA loadings for the five aquaticity dimensions. All five dimensions loaded positively on PC1, which explained 97.0% of total variance; PC2 explained only 2.2%. The near-uniform positive loading confirms that the five dimensions behave as a single coherent gradient rather than opposing axes.

**Supplementary Fig. S2.**
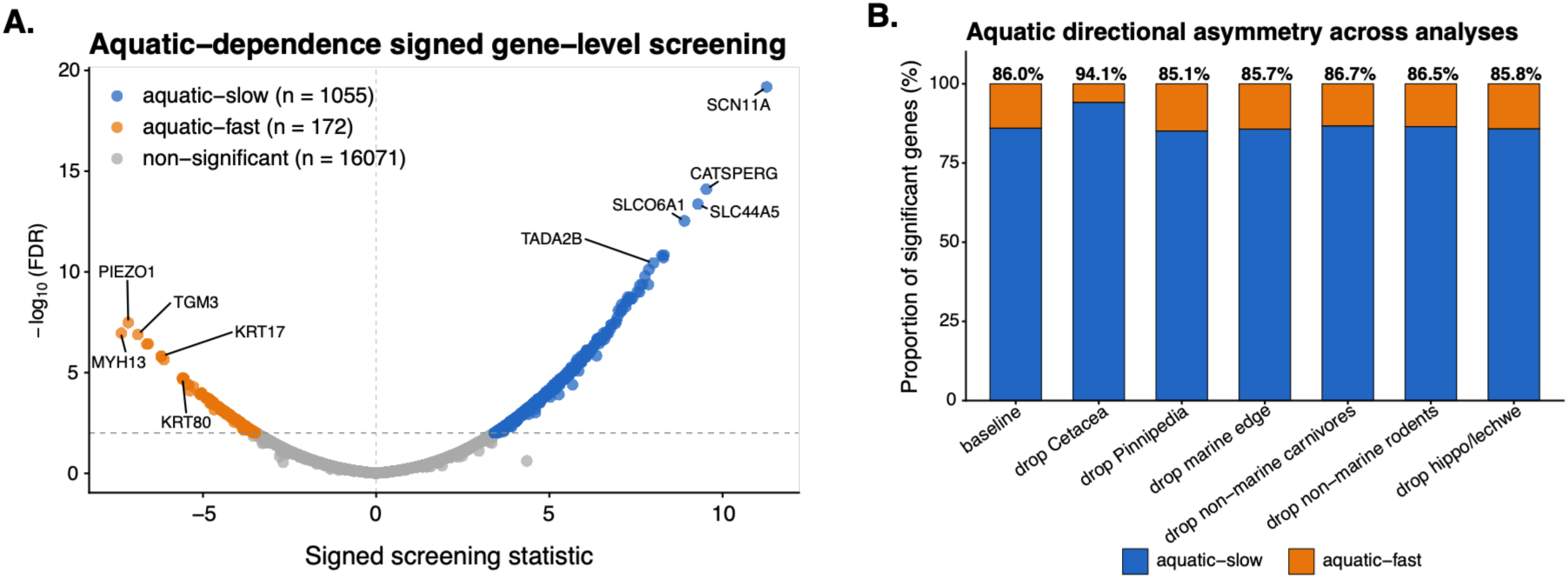
General aquatic dependence shows a parallel slow-direction asymmetry. (A) Signed gene-level screening plot for the aquatic-dependence trait. Of the genes tested, 1,055 were aquatic-slow (blue; lower evolutionary rates on aquatic branches) and 172 were aquatic-fast (orange; higher evolutionary rates), indicating that 86.0% of significant aquatic-associated genes evolved more slowly — a slow-direction bias parallel to the marine screen shown in Fig. 3A. The x-axis shows the signed screening statistic: positive values indicate slower evolution on aquatic-associated branches. (B) This slow-direction dominance was robust across taxon-removal sensitivity analyses (85.1–94.1% slow). Among enriched phenotype and measurement terms, aquatic-slow genes showed a narrower enrichment profile than marine-slow genes, limited to a broad Hematological measurement term; full enrichment results are in Supplementary Table S3.

**Supplementary Fig. S3.**
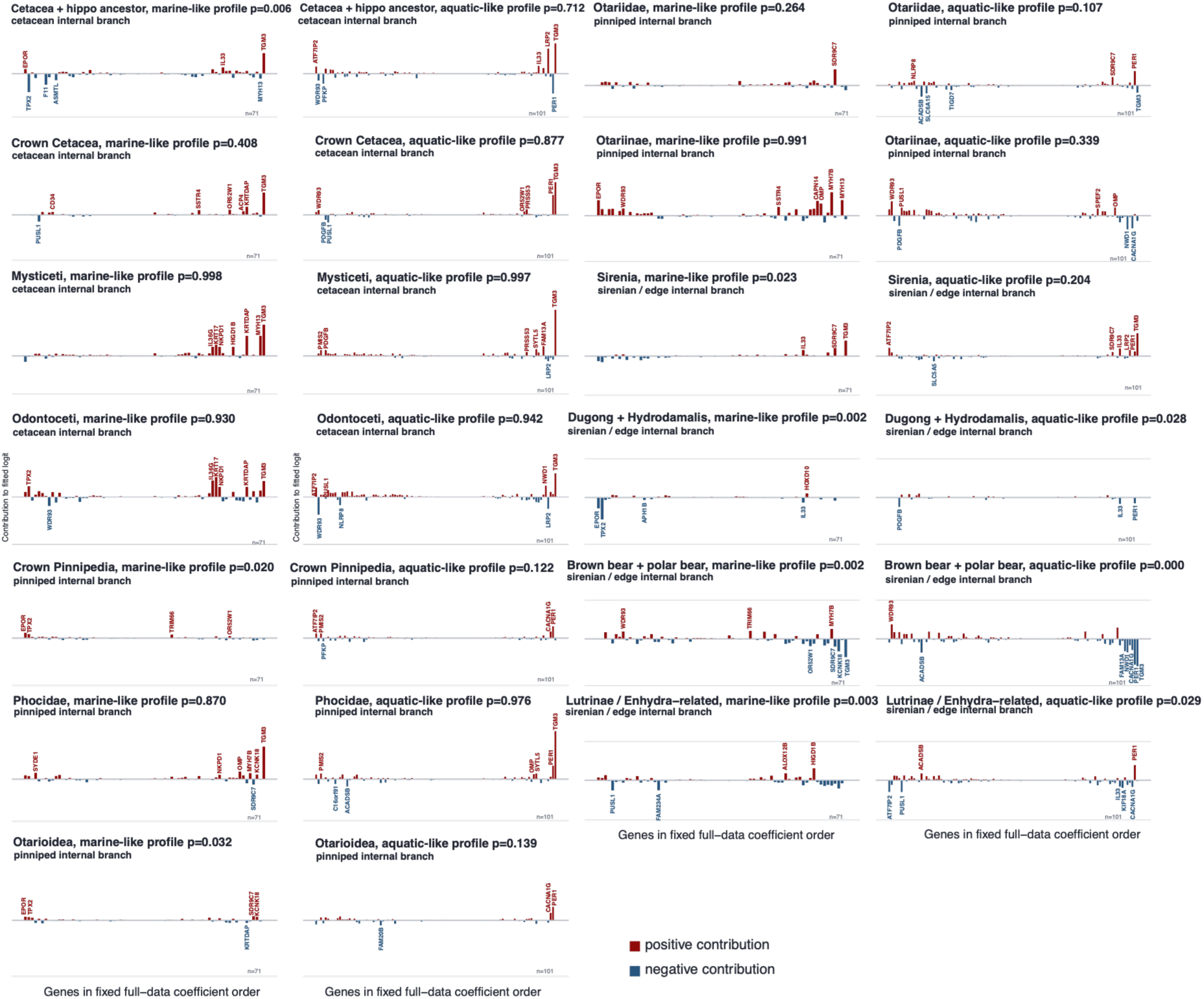
Internal-branch genomic fingerprints reveal how marine-like and aquatic-like profiles were assembled at successive evolutionary stages. Gene-level decompositions for selected ancestral branches, using the same 71 marine-fingerprint genes and 101 aquatic-dependence genes as Fig. 6B,C. Each bar represents the contribution of one gene to the branch’s marine-like or aquatic-like profile score, calculated as scaled GBI × coefficient; the intercept is omitted. Red bars push the profile score toward the marine or aquatic state, whereas blue bars push it away. Gene labels mark the strongest contributors. Branches are grouped by lineage: cetacean internal branches, pinniped internal branches, and sirenian or marine-edge internal branches. Profile scores are rate-based genomic profiles and should not be read as direct habitat assignments. These decompositions complement Fig. 6A by showing not only where ancestral branches fall on the marine and aquatic axes, but which genes contribute to those positions.

**Supplementary Table S1. | Trait definitions and aquaticity scoring across 302 mammals.** Per-species trait assignments including marine-category source, five-dimensional aquaticity scores (foraging medium, locomotion or escape, reproduction or nursery, morpho-physiological specialization, aquatic time budget), summed aquaticity scores, and alternative aquatic coding schemes used for sensitivity analyses. Marine membership was curated as a binary trait from established marine mammal authorities. Semi-aquatic taxa were retained as an intermediate category but were excluded from the primary binary aquatic-dependence endpoint to maintain an unambiguous two-state contrast.

**Supplementary Table S2. | Significant genes from marine and aquatic single-gene screens.** Genes significant at FDR ≤ 0.01 in the branch-level single-gene screening for the marine-binary and aquatic-dependence traits. For each gene, the table reports the screening statistic, significance value, and directional classification as slow (lower evolutionary rates on trait-associated branches) or fast (higher rates). These screening results define the broad trait-associated rate-shift signals underlying the directional asymmetry analyses shown in Fig. 3 and Fig. S2, and provide the candidate-gene pools for downstream LASSO modelling.

**Supplementary Table S3. | Functional enrichment of marine and aquatic screening gene sets.** Functional enrichment results for the directional screening gene sets (marine-slow, marine-fast, aquatic-slow, aquatic-fast), reported across STRING/Monarch phenotype and measurement terms and other STRING-derived annotation categories. The marine-slow enrichment corresponds to the phenotype and measurement summary shown in Fig. 3C. Aquatic-slow enrichment was narrower in the same annotation space, limited among phenotype and measurement terms to a broad Hematological measurement term.

**Supplementary Table S4. | Directional asymmetry and slow-gene overlap across screening analyses.** Summary of slow/fast directional proportions in marine and aquatic single-gene screens across baseline and taxon-removal sensitivity runs. An additional sheet reports marine–aquatic slow-gene overlap and direction-concordance statistics cited in Results Section 2.3: 983 genes were slow-direction in both baseline screens (72.0% of marine-slow genes; 93.2% of aquatic-slow genes), and among 1,138 genes significant in both screens, directional assignments were fully concordant.

**Supplementary Table S5. | Functional annotation of the 148 full-data LASSO predictor genes supporting Fig. 4C**. Gene-level annotation for all 148 genes selected by the marine and aquatic-dependence fingerprints. The table reports predictor partition (shared, marine-specific or aquatic-specific), model coefficients and coefficient direction, compressed display module, recommended function, annotation confidence, evidence notes and Fig. 4C display status. Module assignments are descriptive and are not pathway-enrichment results or evidence of module-level recurrence.

**Supplementary Table S6. | Sensitivity, permutation and predictor-turnover statistics supporting Fig. 5**. Numerical summaries for the lineage-removal cross-validation analyses (Fig. 5A), fold-wise ancestral-state-reconstruction sensitivity checks, the positive-count-matched permutation control for the drop-cetacean marine screen (Fig. 5B), and the predictor-turnover comparisons against comparison-specific candidate-gene null expectations (Fig. 5C). The fold-wise ASR checks masked held-out-genus terminal labels before ancestral-state reconstruction inside each cross-validation fold to verify that the fixed branch-state annotation did not materially affect baseline performance (see Methods).

